# Biomimetic graphitic carbon nitride nanoparticles enable multiscale biomodulation

**DOI:** 10.1101/2024.10.23.619813

**Authors:** Christoph Alexander Müller, Kjeld Kaj Klompmaker, Yuge Zhang, Jing Zhang, Anna Kalatanova, Pengjiu Li, Lingyuan Meng, Jesper Guldsmed Madsen, Thomas Stax Jakobsen, Asbjørn C. Jørgensen, Anne Louise Askou, Yonglun Luo, Lin Lin, Sara Vogt Bleshoy, Georgios Bolis, Ge Huang, Wen Li, Rasmus Schmidt Davidsen, Toke Bek, Nikos S. Hatzakis, Thomas J. Corydon, Henri Leinonen, Bozhi Tian, Mingdong Dong, Menglin Chen

**Affiliations:** Department of Biological and Chemical Engineering, Aarhus University, 8000 Aarhus C, Denmark; Interdisciplinary Nanoscience Center, Aarhus University, 8000 Aarhus C, Denmark; The James Franck Institute, University of Chicago, Chicago, IL, USA; School of Pharmacy, University of Eastern Finland, 70210 Kuopio, Finland; Pritzker School of Molecular Engineering, University of Chicago, Chicago, IL, USA; Department of Ophthalmology, Aarhus University Hospital, 8200 Aarhus N, Denmark; Department of Biomedicine, Aarhus University, 8000 Aarhus C, Denmark; Department of Electrical and Computer Engineering, Aarhus University, 8000 Aarhus C, Denmark; Steno Diabetes Center Aarhus (SDCA), 8200 Aarhus N, Denmark; Department of Chemistry & Nanoscience Center, University of Copenhagen, 1172 Copenhagen, Denmark; Department of Chemistry, University of Chicago, Chicago, IL, USA; The Institute for Biophysical Dynamics, University of Chicago, Chicago, IL, USA

## Abstract

Virtually all organic material on Earth has been produced converting solar energy through photosynthesis in chloroplasts, a sack-like, double membrane organelle in plants and algae, where transmembrane electron transfer occurs from lumen to stroma. Although animals hardly harness the power of photosynthesis, their bioelectrical signals extensively regulate complex electrophysiological behaviors, rendering it a superior target for biomedical innovation. Here, a crude structural mimicry of chloroplast has led us to discover that hollow sphere graphitic carbon nitride nanoparticles (hg-C_3_N_4_ NPs) endowed non-genetic, subcellular and intercellular photo-modulation of various excitable and non-excitable cells, accumulatively achieving modulation at tissue/organ function level. The homogeneous hg-C_3_N_4_ NPs showed responsiveness to light via both photoelectrochemical and photothermal mechanisms. The hg-C_3_N_4_ NPs can be spontaneously internalized with excellent cytocompatibility. Using a focusing laser, the hg-C_3_N_4_ NPs enable intracellular optical stimulation with subcellular resolution, inducing calcium transient release in multiple cells and propagation in primary cardiomyocytes and cardiac fibroblasts. At multicellular scale, optical pacing and synchronization of cardiomyocyte beating is readily achieved by LED. Further, we demonstrate that hg-C_3_N_4_ nanoparticles can be safely delivered into the mouse eye and elicit measurable cortical and behavioral light responses in a subset of animals in a model of advanced retinal degeneration. Finally, application of hg-C_3_N_4_ NPs to porcine retinal tissue *ex vivo* confirmed their modulation capability to directly activate RGCs activity under LED photostimulation. Taken together, these nanostructured biomimetic photocatalytic NPs offer high resolution, leadless optical probing, non-invasive delivery and great biocompatibility, serving as a versatile tool for addressing a range of complex biomedical challenges through subcellular, intercellular and tissue-level photo-modulation across a broad spectrum of scales.

## Introduction

The possibility to electrically sense and stimulate living tissue opens unprecedented potential for therapeutic applications ^1,2^. Particularly, neural and cardiac disease treatment can benefit from materials allowing electrical modulation of their bioelectric activity ^3,4^. As almost all cells are electroactive and respond to electrical signals, electrical stimulation on specific cells needs to be delivered in a precise and controllable manner with high spatiotemporal control ^5^.

Light is characterized by its excellent spatial and temporal stimulation resolution. Optogenetics through genetically introducing photosensitive channel protein, such as Channelrhodopsin-2 found in green algae, has proven to be a versatile tool to control transmembrane and intracellular bioelectric activity by light trigger, with high cell-specific selectivity and spatiotemporality ^6^. However, it requires genetic modifications and concerns immunogenicity of the introduced exogenous transmembrane proteins of often non-mammalian origin ^7^.

Interfaces between biological systems and nanomaterials open an array of possibilities for non-genetic modulation of bioelectric activity with subcellular spatiotemporal control ^8,9^. Semiconductor materials with intriguing photoelectric properties, which can be designed in various shapes and patterns at the nanoscale, are promising candidates to achieve the optically active biointerface ^10,11^. This is especially true for intracellular electrical interfaces, as the natural scale of organelles ranges within tens and several hundreds of nanometers ^12^.

Nanoparticles (NPs) have shown to be able to build tight interfaces with both intra and extracellular membranes ^13,14^. Importantly, light as external power source can trigger electrochemical or photothermal effects at the semiconductor particle/cellular interface acting as a leadless electrophysiological modulator ^15,16^. Leveraging its unique photocatalytic properties, graphitic carbon nitride (g-C_3_N_4_), a polymeric semiconductor, finds applications spanning energy (batteries ^17^, hydrogen evolution ^18^) to biomedicine (biosensors, cancer therapy)^19^. Compared to inorganic nanoparticle photocatalysts (e.g., TiO₂, CoN/CdS), ^20^ g-C_3_N_4_ offers significantly improved visible-light absorption, superior biocompatibility, and greater photostability in physiological environments. Compared to other organic nanoparticles, such as conjugated polymers ^21,22^ or metal organic framework (MOF) ^23^, g-C_3_N_4_ features intrinsic visible-light responsiveness without requiring complex heterojunction engineering or chemical functionalization ^24^. Furthermore, organic semiconductors often suffer from photobleaching, chemical instability, and reduced performance in biological settings, limitations that g-C3N4 materials largely overcome due to their thermal, chemical and photochemical stability. Compared to other photovoltaic systems that typically require large-area implants ^25^, injectability makes NPs a minimally invasive tool for electrophysiological modulation for a broad range of tissues ^14,21^.

Virtually all organic matter on Earth originated from the conversion of solar energy via photosynthesis in chloroplasts, a double membrane organelle in plants and algae, where transmembrane electron transfer proceeds from the lumen to the stroma.^26^ Inspired by this architecture, we utilized hollow-sphere (h)g-C_3_N_4_ nanoparticles as a crude mimicry of chloroplast structure. The hollow morphology improves photoelectronic performance by increasing surface-to-volume ratio, enhancing internal light reflection, and inducing reactive surface strain, while also showing superior cell uptake and intracellular biomodulation capacity ^27^. This biomimetic design enhances electrophysiological modulation at the intracellular and extracellular level, ultimately enabling tissue- and organ-level modulation (Fig. 1a–d). Photocurrent measurements suggest that the photo response of hg-C_3_N_4_ nanoparticles involves both photoelectrochemical and photothermal components, which collectively contribute to the generation of reactive oxygen species (ROS). The hg-C_3_N_4_ NP can be safely internalized with excellent cytocompatibility. The potential of hg-C_3_N_4_ as a tool for intracellular light stimulation with subcellular resolution is demonstrated to induce calcium flux in various excitable and non-excitable cells, allowing cell-cell calcium propagation in primary cardiomyocytes and cardiac fibroblasts. Further, the capacity of hg-C_3_N_4_ NPs for potential cardiac pacing applications was readily achieved by low intensity LED light. At tissue/organ level, the hg-C_3_N_4_ NPs endowed neuromodulation of retinal tissue to reactivate retinal ganglion cell activity after intravitreal injection in blind rd10 mice of autosomal recessive retinitis pigmentosa. This is the first time that biomimetic hg-C_3_N_4_ NPs achieve subcellular, intercellular and tissue-level photo-modulation across a broad spectrum of scales.

**Fig. 1.**
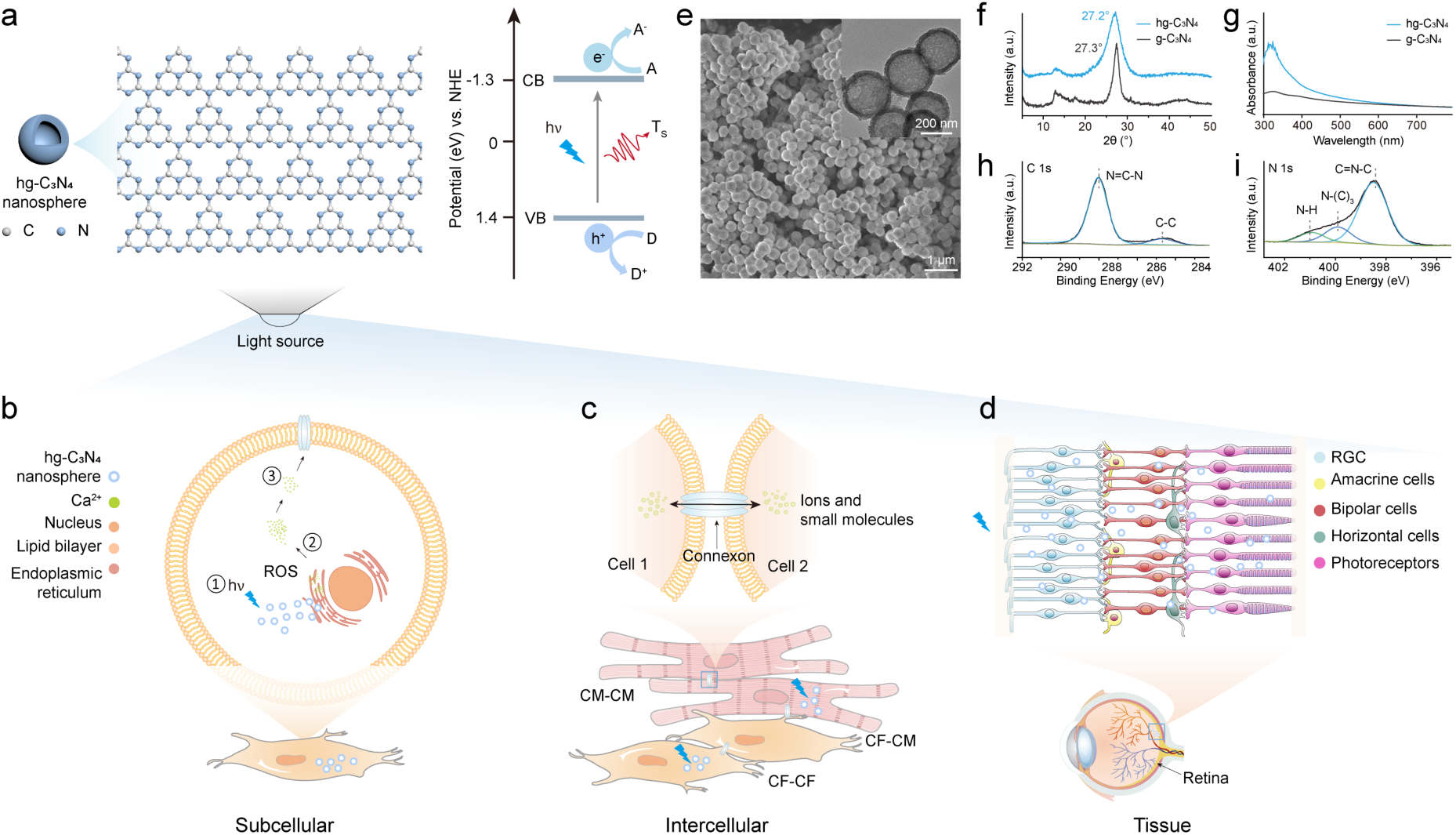
Schematic and characterization of hg-C3N4 NPs for versatile multiscale optical biomodulation. (**a**) The chloroplast mimicking hollow sphere (hg-C3N4) NPs with chemical structure consisting of a covalent triazine network. The bandgap diagram indicates electrons reacting with acceptor (A) compounds and holes reacting with donor (D) compounds, while simultaneously inducing photothermal heating (Ts). (**b**) Proposed mechanism of optical stimulation with subcellular resolution, where photostimulation of hg-C3N4 NPs leads to photofaradaic and photothermal effects along with ROS generation (1), interacting with ryanodine receptor of calcium storing organelle endoplasmic reticulum (2) leads to intracellular calcium flux through the cytosol (3). (**c**) Photostimulation of hg-C3N4 NPs facilitates intercellular electrophysiological interaction of cardiac fibroblast (CF) and cardiomyocytes (CM). (**d**) hg-C3N4 NPs as injectable retina prosthesis stimulating retinal ganglion cells upon photostimulation. (**e**) SEM and TEM (inset) image of hg-C3N4 NPs. (**f**) XRD spectra of hg-C3N4 and bulk g-C3N4. (**g**) UV-vis absorption spectrum of hg-C3N4 and bulk g-C3N4. (**h**) XPS C 1s and (**i**) N 1s spectra of hg-C3N4.

## Results

### Morphological, photoelectrochemical, photothermal and chemical characterizations of the hg-C_3_N_4_ NPs

To achieve hollow spherically shaped hg-C_3_N_4_ NPs, a tailored silica template with subsequent hydrofluoric acid (HF) etching was employed. Scanning electron microscopy (SEM) and transmission electron microscopy (TEM) clearly show the hollow sphere structure of the particles with an inner diameter of around 233 ± 14 nm, an outer diameter of 306 ± 20 nm, and a shell of around 73 ± 14 nm (Fig. 1e).

The XRD spectra (Fig. 1f) shows a (100) reflection peak at ∼13° for both g-C_3_N_4_ and hg-C_3_N_4_, indicating the in-plane periodic arrangement of triazine/heptazine units. The (002) peak of g-C_3_N_4_ at 27.3°, corresponding to interlayer stacking, shifts slightly to 27.2° for hg-C_3_N_4_, indicating an increased interlayer distance from 0.326 nm to 0.328 nm, suggestive of successful structural modification. The great light harvesting properties of hg-C_3_N_4_ NPs could be confirmed by UV-vis spectroscopy, where hg-C_3_N_4_ NPs showed up to 4 times higher light absorbance within a range of 350-600 nm, indicating considerably more absorbance in the visible spectrum compared to bulk g-C_3_N_4_ (Fig. 1g). This effect could be attributed to multiple reflection sites residing in the hollow structure, prolonging the light path length for absorption within the spheres ^27^.

The XPS C 1s spectrum of hg-C_3_N_4_ shows two peaks attributed to N=C–N (288.0 eV) and sp²-hybridized C–C bonds (284.7 eV) (Fig. 1h). The N 1s spectrum reveals three components corresponding to C=N–C (398.5 eV), N-(C)₃ (399.9 eV), and N–H (400.9 eV), confirming the chemical environment of nitrogen atoms in the structure (Fig. 1i). EDX Elemental mapping (Supplementary Fig. S1a) confirms the uniform distribution of C and N elements throughout the hg-C_3_N_4_ sample, supporting its compositional homogeneity.

The stability of nanoparticles directly influences their behavior and efficacy within biological systems, underscoring the importance of studying their stability in various media ^28^. Using Dynamic Light Scattering (DLS) and zeta-potential measurements, the stability and size changes of nanoparticles in various media can be evaluated, providing crucial reference data for the application of nanoparticles. Supplementary Fig. S1b illustrates the stability of nanoparticles in diverse media, including PBS and the cell culture medium, monitored over 12 days. In PBS, the size of hg-C_3_N_4_ NPs in PBS at both day 0 and day 12 is comparable to that of the single NP, indicating excellent suspension stability and minimal aggregation. In parallel, a notable decrease in the zeta potential of NPs after 12 days suggests the formation of an ion corona, which may have enhanced the stability of the hg-C_3_N_4_ NPs suspension. In DMEM with 10% FBS, the formation of the protein corona may have shielded a portion of the charges on the particles, leading to minor changes in the zeta potential after 12 days. The observed increase in particle size can be attributed to protein adsorption onto the particles, leading to the formation of a protein corona ^29^. The formation of protein corona can lower the surface energy of hg-C_3_N_4_ NPs and promote their dispersion in biological fluids ^30^.

Leadless bioelectric stimulation using photo-responsive particles relies on photoelectrochemical or photothermal effects, both of which have been shown to induce membrane depolarization in adjacent cells ^15,31^ (Fig. 2a). The relationship of photocurrent trace amplitude and applied holding currents I_0_ provides insight to the nature of the generated photocurrents. A patch-clamp electrophysiology apparatus with integrated photocurrent measurement set-up was employed to measure the photocurrent (Fig. 2b). As photothermal effects go along with temperature increase, the increased ion mobility leads to reduced pipette resistance and subsequently increased current, directly proportional to the amperage of I_0_. On the other hand, photoelectrochemical processes, like capacitive and faradaic currents, are independent from the applied holding current ^32^. The non-zero intercepts in Fig. 2c-d along with the slope of photocurrent amplitudes across holding currents in Fig. 2e, indicate the presence of anodic currents and a transient temperature increase of approximately ΔT = 0.2325 °C. Together, these findings suggest the coexistence of photoelectrochemical and photothermal effects. While a less pronounced anodic current (–5 pA at I₀ = 0 nA) was observed under a 10 ms light pulse (Fig. 2c), the sustained negative current (–30 pA at I₀ = 0 nA) throughout the 100 ms pulse (Fig. 2d) serves as evidence of an anodic faradaic response, in contrast to capacitive currents characterized by rapid charge–discharge dynamics. These results confirm the coexistence of photothermal and faradaic effects under illumination ^32^, consistent with observations in other nanotransducers for photostimulation ^33^.

**Fig. 2.**
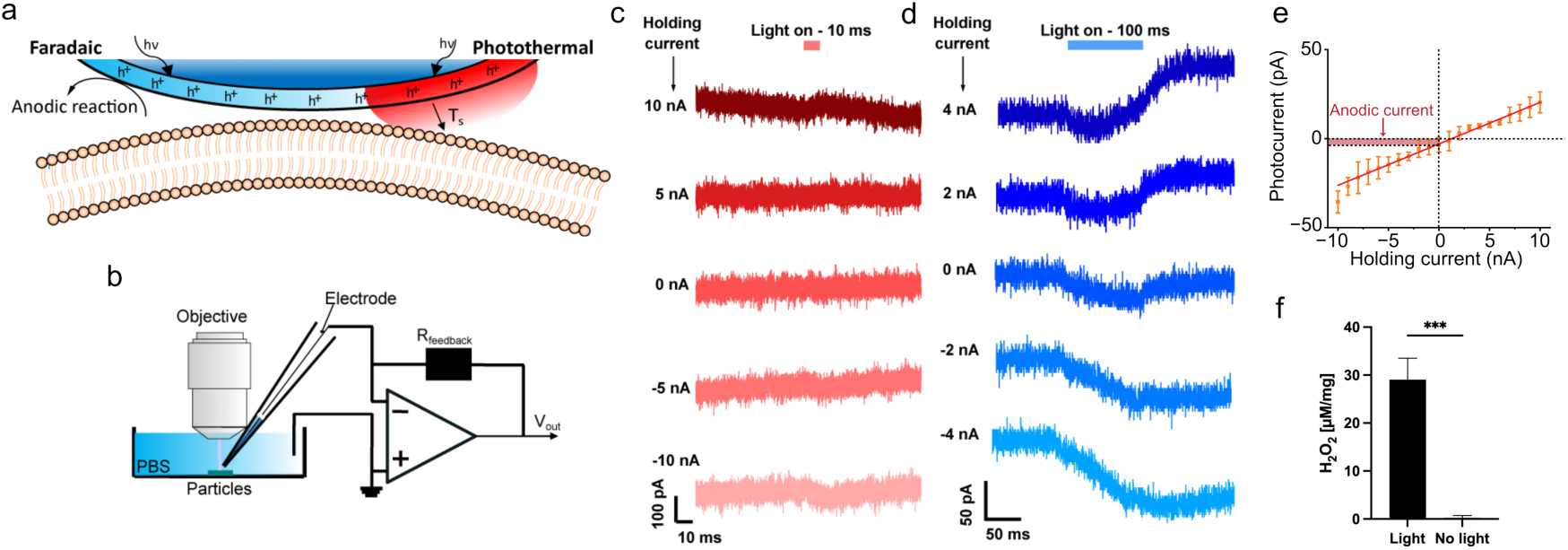
Photocurrent characterization. (**a**) Scheme of hg-C3N4 NPs stimulation mechanism based on anodic faradaic photocurrent and photothermal effect Ts, light-induced heating; hν, photon energy. (**b**) Illustration of patch-clamp photo-response measurement set-up used to measure photocurrent. (**c-d**) Photocurrent traces measured for a range of holding currents (I₀) under 625 nm LED illumination at 45 mW/mm² intensity, with (c) 10 ms and (d) 100 ms pulse durations. (**e**) Photocurrent amplitudes for a range of holding holding currents. Data are shown as means ± SD (n = 3). (**f**) Quantification of hg-C3N4 NPs produced H2O2 under visible-light LED illumination (75 mW/cm^2^). Data (n = 7) is shown as means ± SD. Statistical analysis was perfomed using unpaired two-tailed t-test where p<0.0001 is denoted as ***

Since faradic photocurrents go along with redox reactions at the semiconductor/electrolyte interface, the generation of ROS is probable. In addition, photothermal effect has also been reported for ROS generaton ^34,35^. The quantification of H_2_O_2_ (Fig. 2f, Supplementary Fig. S2a) using a p-hydroxyphenylacetic acid (pHPA)/horseradish peroxidase (HRP) assay revealed a clear dose-dependent increase in H₂O₂ production. Under blue light irradiation (450 nm, 75 mW/cm²) for 30 minutes, approximately 30 µM of H₂O₂ was generated per mg of hg-C_3_N_4_, while no H₂O₂ was detected in the absence of light.

Using electron spin resonance (ESR) ^36^, a more direct and chemically specific method compared to fluorescent probes, the presence of intermediate ROS radicals, superoxide anion •O_2_-, hydroxyl radicals •OH, and singlet oxygen ^1^O_2_, was detected (Supplementary Fig. S2b-d). This suggests an electrochemical process that produces H_2_O_2_ through the oxygen reduction pathway, which reduces O_2_ to •O_2_-, or through the water oxidation reaction, which oxidizes H_2_O to •OH, or both ^37^. Due to the inferior oxidation capability of hg-C_3_N_4_, formation of hydroxyl radicals •OH is most likely attributed to H_2_O_2_ photolysis into •OH, or to Fenton-like reaction promoted by photothermal effects ^38^. Singlet oxygen (^1^O_2_) can be formed by the deactivation of O_2_•- on the photocatalyst surface ^39^. Therefore, the reduction of oxygen to superoxide anion •O_2_- is the major contributor for H_2_O_2_ formation through subsequent reaction with H^+^.

To verify whether this extracellular ROS production translated into intracellular oxidative signaling, NIH/3T3 fibroblasts, used here as a general cellular model, were incubated with the redox-sensitive probe 2’,7’-dichlorodihydrofluorescein diacetate (DCFH-DA). Upon blue light exposure (10 mW/cm²) for 10 minutes, cells treated with hg-C_3_N_4_ showed a significant increase in fluorescence relative to controls, indicating intracellular redox activity (Supplementary Fig. S3).

### Particle internalization and cytocompatibility

For responsive nanomaterials intended to interface with excitable cells, cellular internalization is a critical property, as it enables interaction with intracellular compartments and signaling pathways at the subcellular level. We therefore investigated whether hg-C_3_N_4_ nanoparticles are efficiently internalized by mammalian cells and their intracellular trafficking properties.

NIH/3T3 mouse embryonic fibroblasts were used as a general and well-established cellular model to study nanoparticle uptake and intracellular trafficking ^40^. To characterize the internalization of hg-C_3_N_4_ NPs, the inherent blue fluorescence of hg-C_3_N_4_ was used to track the particles over time, using our toolboxes ^41–43^. Single NIH/3T3 fibroblasts labeled with a membrane marker were continuously imaged (Fig. 3a, Supplementary Fig. S4a) and the increase of hg-C_3_N_4_ NPs over time showed their successful, albeit slow, internalization (Fig. 3a, c).

**Fig. 3:**
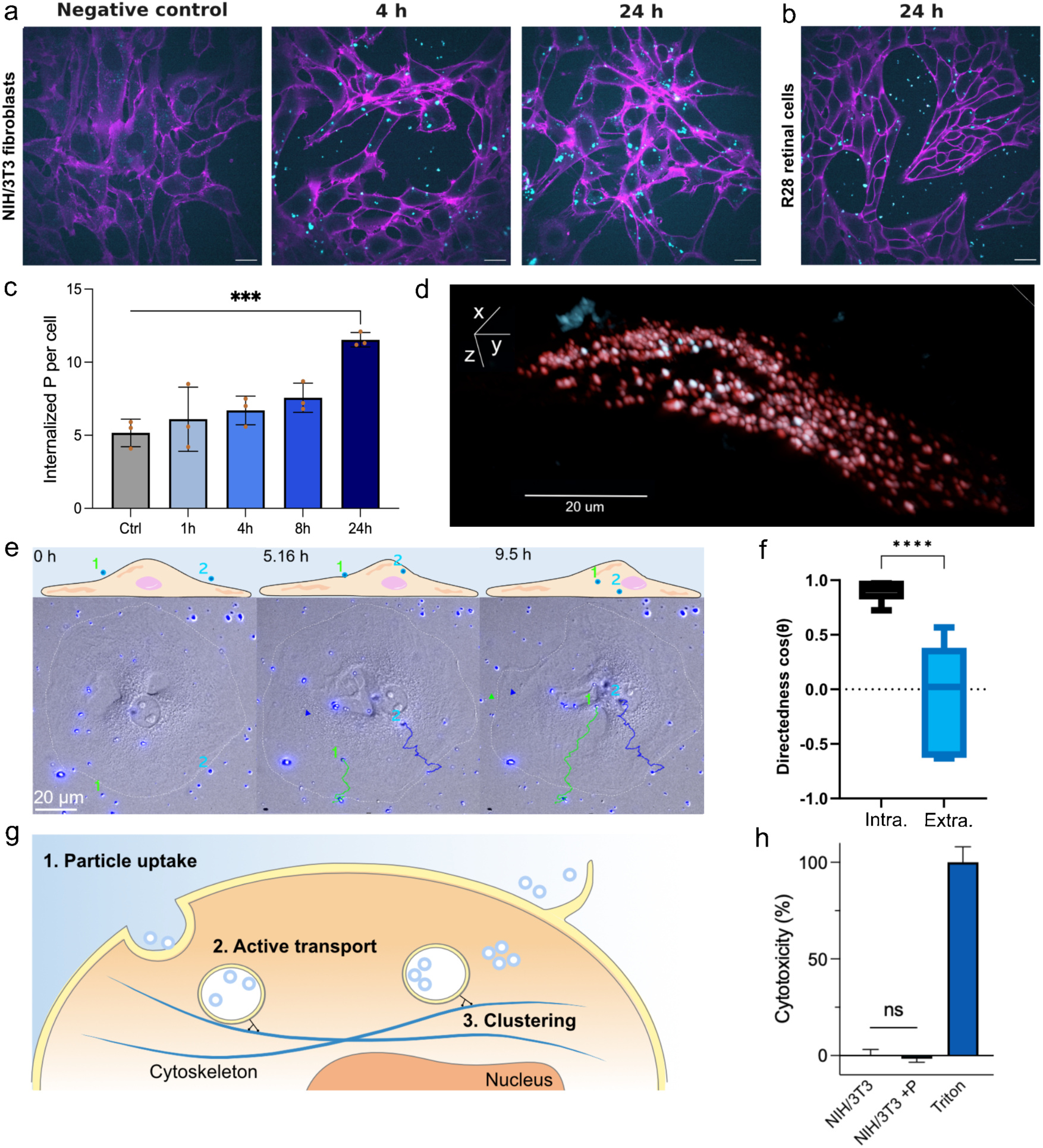
Internalization and cytocompatibility of hg-C3N4 NPs. (**a**) Live cell images of the time-dependent internalization of hg-C3N4 NPs (cyan) in NIH/3T3 fibroblasts (plasma membrane labeled in magenta), acquired on spinning-disk confocal microscope. Scale bar is 20 µm. (**b**) Representative live-cell image of hg-C3N4 NP uptake in R28 retina cells acquired in the same manner as (a). Scale bar is 20 µm. (**c**) Quantification of internalization of hg-C3N4 NPs in NIH/3T3 fibroblasts (n > 500 cells). Data are shown as mean ± SEM, with individual biological replicates shown in orange. Statistical analysis was performed using one-way ANOVA where p<0.001 is denoted as ***. (**d**) 3D view acquired by lattice light-sheet microscopy of a single NIH/3T3 cell showing internalized hg-C_3_N_4_ NPs (cyan) and lysosomes (red), with colocalization shown in white. Scale bar is 20 µm. (**e**) Tracking of two representative hg-C3N4 NPs internalized by a NIH/3T3 cell over time. (**f**) Quantification of directedness for internalized (intra.) and non-internalized (extra.) particles (n = 8). Statistical analysis was performed using unpaired two-tailed t-test where p<0.0001 is denoted as ****. (**g**) Schematic of the proposed internalization and intracellular trafficking mechanism. (**h**) LDH cytotoxicity assay of hg-C3N4 NPs in NIH/3T3 fibroblasts. Data (n = 6) are shown as mean ± SD

To assess whether this uptake behavior extends beyond fibroblasts, we additionally examined hg-C_3_N_4_ internalization in R28 retinal cells. Similar particle uptake was observed qualitatively (Fig. 3b, Supplementary Fig. S4b), indicating that hg-C_3_N_4_ NPs can be internalized by distinct mammalian cell types. Definitive intracellular localization was confirmed using volumetric imaging in NIH/3T3 cells, which further showed the hg-C_3_N_4_ NPs to colocalize with lysosomes after prolonged exposure (Fig. 3d, Supplementary Video S1).

Tracking of the particles revealed a clear directionality from the extracellular space toward the perinuclear space (Fig. 3e-f, Supplementary Fig. S4c, Supplementary Video S2). Directedness was used as a quantitative indicator to distinguish internalized particles from non-internalized ones based on their loss of random mobility (Fig. 3f). Internalized particles exhibited a median cos(θ) value of 0.91, corresponding to movement within ±25° toward the nucleus, whereas non-internalized particles showed broadly distributed cos(θ) indicative of random motion ^44^. This reduction in random mobility provides a strong quantitative indicator of successful particle internalization.

The factors that influence the mechanism of internalization are manifold, with particle size and shape being among the most important particle properties ^45,46^. To investigate the uptake pathways involved, we performed a chemical inhibitor assay ^47^. While chloroquine showed a slight decrease across all biological replicates, no single inhibitor was able to significantly reduce internalization (Supplementary Fig. S4d, Supplementary Table S1). Combined with the trafficking behavior observed above, these findings suggest that the hg-C_3_N_4_ NPs are internalized through multiple pathways, including both phagocytosis- and endocytosis-driven mechanisms (Fig. 3g) ^48,49^, followed by active trafficking within endosomal compartments toward the perinuclear region ^50^.

To assess the cytocompatibility of hg-C_3_N_4_ NPs, we evaluated their effects on cellular viability using complementary in vitro assays. First, lactate dehydrogenase (LDH) release was used as a basal cytotoxicity assay in NIH/3T3 cells incubated with the NPs. This assay did not reveal any considerable cytotoxic effects of hg-C_3_N_4_ NPs in NIH/3T3 cells (Fig. 3h).

As photostimulation of hg-C_3_N_4_ NPs could potentially induce phototoxic effects, we next examined the impact of high-dose blue LED light exposure (450 nm, 75 mW/cm^2^, 1 Hz, 100 ms pulse width, 10 minutes bidaily). Live/dead staining of NIH/3T3 fibroblasts following acute light exposure did not reveal detectable cytotoxic effects compared to non-illuminated controls (Supplementary Fig. S5a-b). Light exposure led to a transient reduction in proliferation at day 3; however, proliferation recovered by day 8 in the presence of hg-C_3_N_4_ NPs (Supplementary Fig. S5c). As high-intensity blue light has previously been reported to negatively affect cellular viability ^51^, all subsequent experiments were performed using lower light intensities (10-30 mW/cm^2^).

To exclude potential confounding effects of cell division, acute LDH assays and live/dead staining following repeated photostimulation over 7 days were additionally performed in non-dividing human induced pluripotent stem cell-derived cardiomyocytes (iPSC-CMs), in which minimal toxicity was observed (Supplementary Fig. S6).

### Biological modulation at subcellular and cellular level

#### Subcellular photostimulation of non-excitable cells

The ability of the particles to induce light-triggered intracellular bioelectric activity with subcellular resolution was investigated by monitoring calcium flux under focused laser beam stimulation of single internalized particles. hg-C_3_N_4_ NPs showed strong fluorescence in the green channel, enabling simplified identification of internalized hg-C_3_N_4_ NPs. A focused 473 nm (8.8 mW/µm^2^) laser beam with a pulse width of 200 ms and spot size of 3 µm with subcellular resolution (Supplementary Fig. S7) was employed to stimulate the particles. The laser pulse alone could not provoke any change in calcium dynamics in NIH/3T3 fibroblast cells (Supplementary Video S3). When particles were present, photostimulation led to an immediate calcium flux through the cytosol originating from the stimulated particle location (Fig. 4a, Supplementary Video S4–5), as evidenced by the vector direction graphic. For NIH/3T3 cells, the average calcium flux velocity was 2.4 ± 3.8 µm/s.

**Fig. 4.**
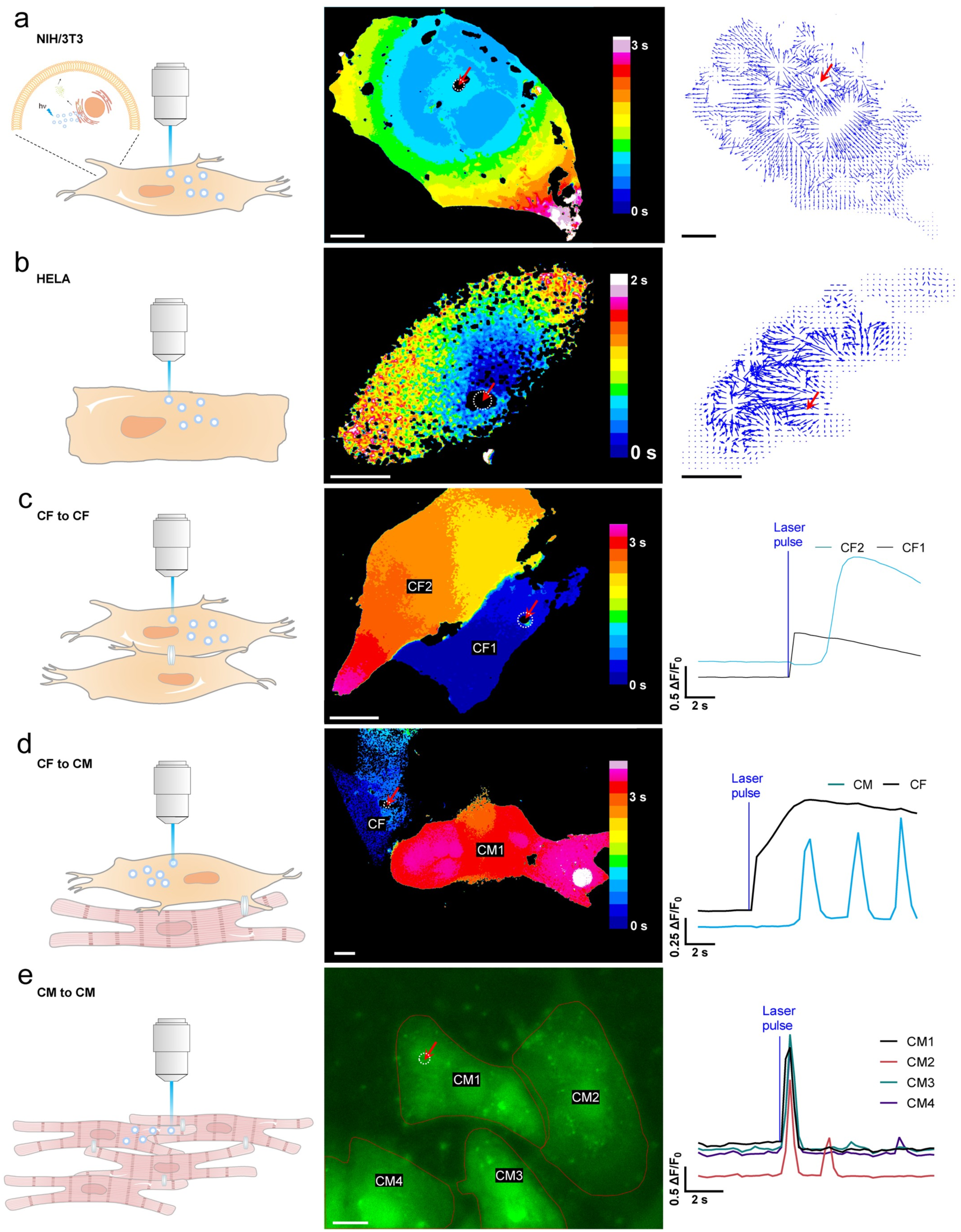
Subcellular and intercellular electrical interrogation. Intracellular electrical interrogation of NIH/3T3 fibroblasts **(a)** and HELA cell (**b**) with corresponding isochronal and vector map. Inset in (a) shows the proposed calcium release mechanism based on calcium storing ER. **(c)** Intercellular calcium propagation with corresponding isochronal maps or calcium staining with ΔF/F0 plots between CFs, CF and CM (**d**), CM and CM (**e**). Scale bars are 10 µm. Red arrow indicates location of 200 ms laser pulse, 473 nm, spot size 3 µm, 8.8 mW/µm^2^.

The versatility of hg-C_3_N_4_ NPs to study intracellular bioelectric activity of non-excitable cells was also demonstrated by HELA cells (Fig. 4b). A single 200 ms laser pulse led to an immediate intracellular calcium flux with a prolonged increase in intracellular calcium, evident by the sustained increase in fluorescence intensity (Supplementary Video S6–7). The average velocity of the calcium flux in HELA cells was determined to be 1.3 ± 2.7 µm/s. Extended recording up to 50 s (Supplementary Fig. S8) revealed detailed kinetics, including the time-to-peak (ToP ≈ 1.9 s) and decay constant (τ ≈ 27.7 s), demonstrating that the cytosolic Ca²⁺ signal persists well beyond the original 8 s window. These longer observations confirm the stability of the response and provide quantitative insight into the elevation and recovery dynamics.

#### Photostimulation of excitable cells and intracellular signal propagation

The application of hg-C_3_N_4_ NPs to study intracellular bioelectric activity of excitable cells was then demonstrated by cardiomyocytes (CM), which play a key role in heart tissue muscle contraction. We tested the ability of hg-C_3_N_4_ NPs to electrically modulate primary rat CMs and primary rat cardiac fibroblasts (CFs).

Cell-cell signal propagation was observed not only between the CMs, but also CFs. CFs express a plethora of ion channels, mainly potassium (K^+^) channels and mechanosensitive channels, that contribute to their resting membrane potential ^52–55^. Indeed, a stimulated primary CF propagated the calcium transient to an adjacent CF with a velocity of 8.8 ± 16.2 µm/s (Fig. 4c, Supplementary Video S8). Direct coupling of cardiomyocytes and fibroblasts via connexin-based gap junctions is supported by compelling evidence ^56–58^. Stimulation of a primary CF adjacent to CM led to a similar observation, when the intracellular calcium flux of the fibroblast reached the outer cell membrane with 4.4 ± 8.5 µm/s, the adjacent CM started with continues beating (Fig. 4d, Supplementary Video S9). hg-C_3_N_4_ NPs associated with CMs were directly stimulated, resulting in immediate intracellular calcium flux and beating (Fig. 4e, Supplementary Video S10). Intriguingly, the calcium flux of the stimulated cell led to calcium flux and beating of three adjacent coupled CMs. This mechanism of signal propagation is typical for coupled CMs and relies on interconnected connexin gap junctions, enabling synchronous beating essential for the physiological function of the heart ^57^.

To further clarify the spectral dependence of cell activation, we compared the efficacy of cell activation using lasers with wavelengths both above and below 500 nm under identical conditions (∼1 mW, 100 ms pulse duration). When irradiated with a laser of >500 nm (635 nm laser), CM cells showed negligible changes in intracellular calcium signaling compared to the pronounced response observed with 473 nm illumination (Supplementary Fig. S7 and Supplementary Video S11, S12). This finding is consistent with the absorbance spectrum of our hg-C_3_N_4_ nanoparticles, in which the absorbance is significantly lower than 500 nm, thus reducing the photoelectrochemical activity required for effective cellular stimulation.

We also demonstrated that the photocurrent generated by hg-C_3_N_4_-coated substrates under repeated 473 nm laser pulses is highly stable across multiple on–off cycles (Supplementary Video S13).

#### Molecular pathway of calcium release

To rigorously determine the molecular pathway underlying the observed calcium elevation, we employed a series of pharmacological inhibitors that target specific calcium signaling components. These experiments allowed us to isolate the functional contributions of ROS, endoplasmic reticulum (ER) calcium channels, and membrane ion channels, providing mechanistic clarity.

Pretreatment with 500 µM N-acetylcysteine (NAC), a well-characterized ROS scavenger, for 1 hour prior to laser stimulation completely suppressed the calcium response (Supplementary Fig. S9 and Supplementary Video S14). This result demonstrates that intracellular ROS generation is essential for triggering calcium release, given our observation of H₂O₂ generation by hg-C_3_N_4_.

Both photothermal and photoelectrochemical stimulation mechanisms have shown to be able to elicit action potentials in neurons and have been used to modulate cardiac tissue ^9,59–62^. To test whether plasma membrane thermosensitive ion channels were involved, we used 10 µM Ruthenium Red (RR), a non-specific blocker of transient receptor potential (TRP) channels. Calcium transients remained intact in the presence of RR (Supplementary Fig. S9 and Supplementary Video S15), excluding the contribution of thermosensitive TRPV/TRPM family channels and arguing against a photothermal membrane-gating mechanism.

Further mechanistic studies were conducted to determine how hg-C_3_N_4_ NPs activate calcium release from the ER. The mammalian ER predominantly expresses two tetrameric calcium channels: the inositol-triphosphate receptor (IP_3_R) and the ryanodine receptor (RyR) ^63,64^. To test whether RyR are directly responsible for calcium release, we pretreated cells with 25 µM ryanodine, a RyR inhibitor, for 10 minutes. Under these conditions, no significant calcium flux was observed after laser stimulation (Supplementary Fig. S10 and Supplementary Video S16). This confirms that the RyRs on the ER membrane are the key mediators of intracellular calcium release in our system. This is consistent with previous reports of redox-sensitive RyR-gating by ROS such as H₂O₂.

We further examined the role of IP₃Rs by treating cells with 200 µM 2-APB, an inhibitor of IP₃-mediated ER calcium release. As shown in Supplementary Fig. S10 and Supplementary Video S17-S18, 2-APB-treated cells still exhibited robust calcium transients following stimulation, indicating that IP₃Rs are not required for the laser-induced Ca²⁺ release. This distinguishes the response mechanism from canonical Gq/PLC/IP₃ signaling pathways. Moreover, the presence of continued spontaneous calcium oscillations in the presence of 2-APB and the absence of global fluorescence loss argues against non-specific ER rupture or membrane damage as the source of calcium release.

Finally, to definitively test whether extracellular calcium influx was required in excitable cells, we conducted stimulation experiments in supplemented Ca²⁺-free HBSS (Supplementary Fig. S11 and Supplementary Video S19). The calcium response remained robust, confirming that the source of Ca²⁺ is entirely intracellular and independent of store-operated calcium entry (SOCE) or voltage-gated calcium channels. In contrast, when examined in non-excitable cells such as HeLa, light-evoked responses were diminished in Ca^2+^-free media (Supplementary Fig. S12), suggesting a contribution from extracellular Ca^2+^ ^65^. Further investigations will be needed to delineate the precise mechanisms in non-excitable systems, as the present study primarily focuses on excitable cell responses.

Together, these inhibitor-based experiments comprehensively support a mechanism in which photoelectrochemical and photothermal effects collectively generated ROS activates intracellular RyRs, leading to ER calcium release. In excitable cells, the process is independent of IP₃R signaling, extracellular calcium influx, TRP channels, and thermal effects, reinforcing intracellular ROS-triggered ER calcium release as the central mechanism of stimulation in this system.

#### Interval training of cardiomyocytes

The injectability of NPs makes them an attractive non-invasive alternative to conventional electronic pacemaker therapy. As the stimulation of single cardiomyocytes via laser has proven feasible, we next wanted to apply interval training to many cardiomyocytes simultaneously to synchronize beating and align to common beating frequency using low intensity LED light source (10 mW/cm^2^, λ=450 nm). When HL-1 cardiomyocytes were cultured together with hg-C_3_N_4_ NPs and LED photostimulation with a frequency of 1 Hz and 100 ms pulse width was applied for 10 min, the beating frequency changed from an irregular non-synchronized pattern between 0.5 and 2 Hz to a more synchronized pattern of 2 Hz (Fig. 5a, b). This effect was even more pronounced after 30 min of stimulation. Further, the calcium flux appeared to travel as a directional synchronized wave through the cardiomyocyte cell sheet, when both hg-C_3_N_4_ NPs and photostimulation were applied (Fig. 5c, Supplementary Video S20-22). Cardiomyocytes cultured without hg-C_3_N_4_ NPs remained in a state of unsynchronized beating behavior. Light stimulation alone led to an increase in beating frequency around 1.5 Hz, however, without the desired synchronization and directionality of the calcium wave (Supplementary Fig. S13). Interestingly cardiomyocyte beating was not synchronized to the stimulation frequency of 1 Hz, but to 2 Hz. This might be attributed to the fact the natural heart rate of mice is much higher than 1 Hz, and to our knowledge no decrease of beating frequency has yet been achieved in pacing experiments ^59,61,66^. This could mean that cardiomyocytes were trained at 1 Hz at their every second beat to achieve a final beating rate of 2 Hz.

**Fig. 5.**
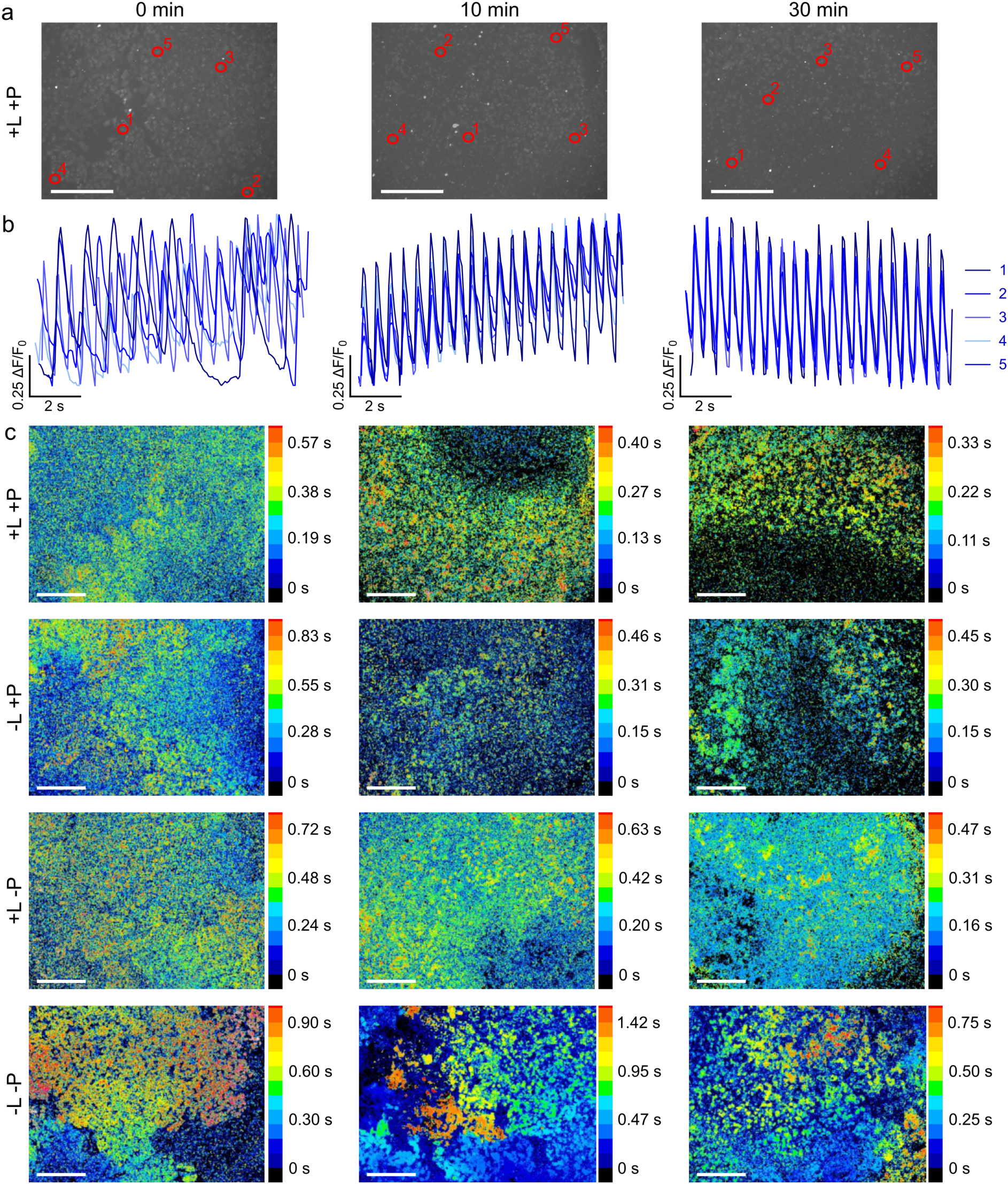
Pacing of cardiomyocyte via photostimulation of hg-C3N4 NPs. (**a**) HL-1 cells stained with Fluo-4 calcium stain incubated with 2.0 µg/cm^2^ hg-C3N4 NPs (+P) and photostimulation (+L) at 0 min, 10 min, and 30 min. Red circles indicate ROIs for ΔF/F0 plots. (**b**) ΔF/F0 plots of 5 different ROIs showing beating frequency of photostimulated HL-1 cells. (**c**) Isochronal maps of Hl-1 cells incubated with or without hg-C3N4 NPs (+P; -P) and with or without photostimulation (+L; -L). Light source was 10 mW/cm^2^ of 450 nm with pulse width of 100 ms and pulsed at 1 Hz frequency. Scale bars are 275 µm.

Furthermore, light-induced biomodulation of human cardiac cells was achieved, as evidenced by calcium imaging of nanoparticle-treated iPSC-CMs, which showed a consistent increase and stabilization at the targeted 1 Hz beating frequency under blue-light stimulation (450 nm, 30 mW cm⁻², 1 Hz, 50 ms pulses) (Supplementary Fig. S14 and Supplementary Video S23-S24).

#### Propagation Mechanism

ATP secretion and gap junctions are known to be responsible for the propagation of intercellular Ca^2+^ waves ^67^. Released ATP can bind to purinergic receptors on adjacent cells, initiating calcium release from internal stores. Gap junctions enable the direct passage of calcium ions and other signaling molecules between cells. Together or separately, these mechanisms coordinate cellular activity.

To elucidate the contribution of these pathways in our system, HL-1 cells were pretreated with either apyrase, an ATP-hydrolyzing enzyme, or carbenoxolone (CBX), a gap-junction inhibitor, for 15 minutes prior to a round of light stimulation (450 nm, 10 mW/cm2, 1 HZ, 100 ms pulses, 10 minutes). Apyrase showed no significant effect on Ca^2+^ wave dynamics, while CBX significantly reduced wave propagation and disrupted its directionality (Supplementary Fig. S15 and Supplementary Video S25-S27). These findings suggest that gap junctions play a dominant role in mediating Ca^2+^ wave propagation in our system.

### Therapeutic biomodulation

The eye is a remarkable organ, and retinal tissue is an essential part of this system, playing a critical role in converting energy from photons of visible wavelengths into electrical signals, preprocessing the visual signals, and then relaying these signals to the brain for further interpretation. This complex retinal function relies on the interplay of different types of neurons within the tissue. Among these, retinal ganglion cells (RGCs) receive input from sensory photoreceptors via retinal interneurons (bipolar cells) and relay this information to the visual areas of the brain. Bioelectronics have emerged as a promising approach for restoring vision, especially if the device is responsive to light ^68^. Here, we further explore the potential of hg-C_3_N_4_ NPs as an injectable retinal prosthesis using living mice as well as porcine retina *ex vivo*.

Importantly, building on the physicochemical stability analysis described previously, we evaluated the long-term stability and biodegradation profile of these nanoparticles to confirm their suitability in vivo. After 30 days of incubation in simulated physiological conditions (pH 7.4 and 7.0), the nanoparticles retained their structural integrity, shape, and dispersion characteristics (Supplementary Fig. S16), thus reducing concerns about degradation-driven toxicity or functional loss.

Encouraged by these findings, we next tested the feasibility of safe delivery to the retina by assessing biodistribution and retinal integrity following intravitreal delivery to wild-type (WT) C57BL/6JRj mice (Fig. 6a). The intravitreal injection of hg-C_3_N_4_ NPs into mouse eyes resulted in the visible settlement of particle aggregates on the retinal surface immediately adjacent to the RGCs and their axons as observed in fundus images and optical coherence tomography (OCT) scans 14 days post-injection (Fig. 6b). OCT imaging demonstrated preserved retinal layer integrity (Fig. 6b-c), where neither total retinal thickness (p = 0.57) nor individual layers thicknesses differed significantly between baseline and day 14 post-injection. The hg-C_3_N_4_ NPs could be clearly identified on top of the RGCs using fluorescence imaging of retinal cross sections (Fig. 6d-f).

**Fig. 6.**
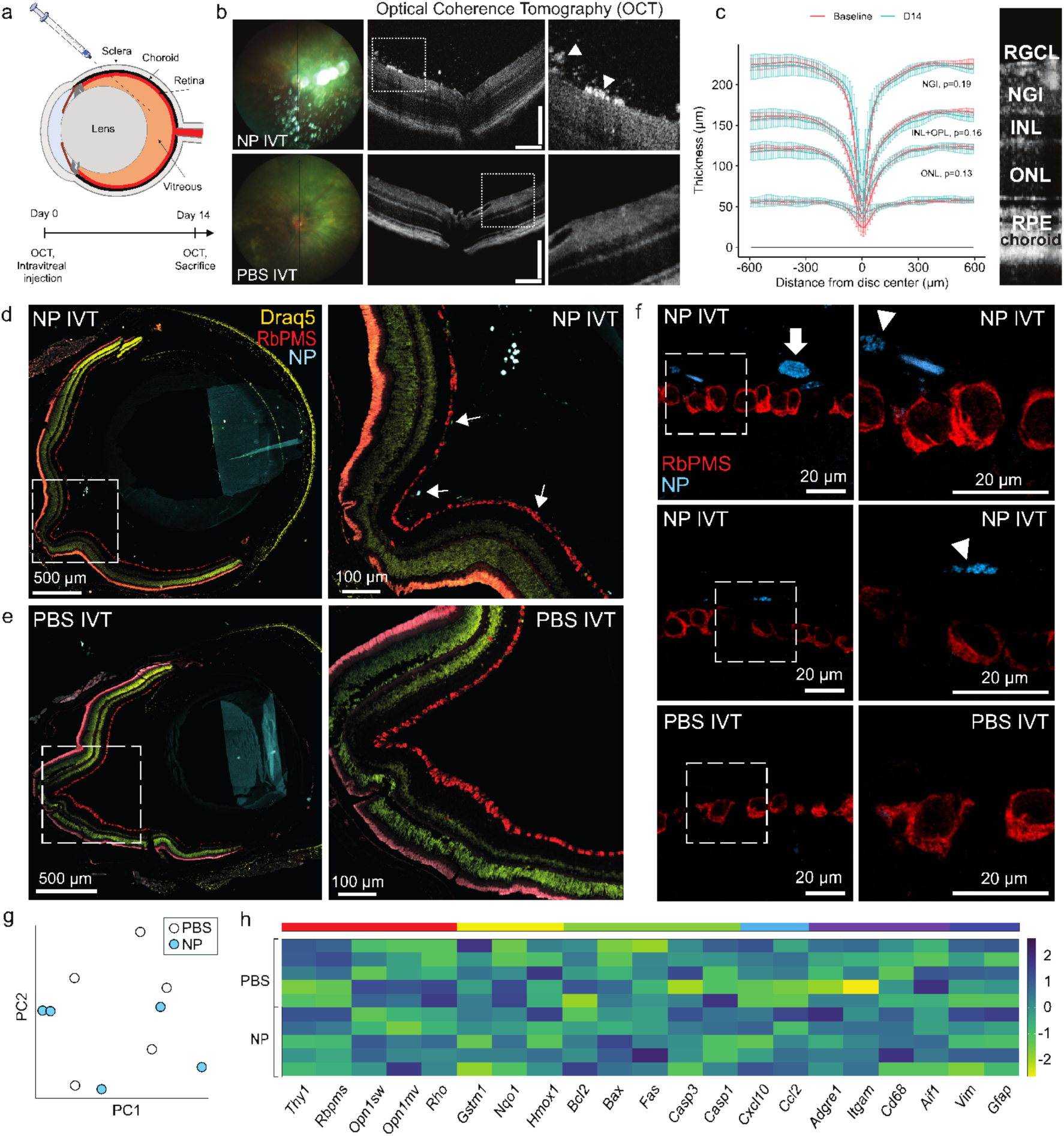
Intravitreal delivery of hg-C3N4 NPs preserves retinal structure and gene expression. (**a**) Schematic drawing of the intravitreal injection procedure and experimental timeline. (**b**) Representative fundus images and optical coherence tomography (OCT) B-scans from eyes intravitreally injected with hg-C3N4 NPs (NP IVT) or buffer (PBS IVT). The integrity of retinal layers is preserved and hyperreflective spots corresponding to aggregated NPs (white arrowheads) can be observed in the vitreous and on the retinal surface. Scale bars are 200 µm. (**c**) Position of retinal borders in the NP IVT eyes at baseline and D14 (mean ± SD) with corresponding comparisons of the mean thickness across individual retinal layers (n = 6). NGI = combined thickness of retinal nerve fiber layer, ganglion cell layer and inner plexiform layer; INL = inner nuclear layer; OPL = outer plexiform layer; ONL = outer nuclear layer. Paired two-tailed t-test was used. (**d-e**) Widefield fluorescent images (20x) from the mice shown in (b) labeled with the RGC marker RbPMS (red) and counterstained with Draq5 (yellow). The blue fluorescent NPs (white arrowheads) are attached to vitreous strands and the retinal surface. The orange signal detected in the outer segments is attributable to autofluorescence. Scale bars are 500 µm. For the inserts the scale bars are 100 µm. (**f**) Confocal microscopy images (63x) of retinal cross sections labeled with the RGC marker RbPMS (red). Blue fluorescent NPs (white arrowheads) are seen on the inner retinal surface near the RGCs. In some areas, NPs were engulfed by rounded cells, presumably hyalocytes (white arrow). Scale bars are 20 µm. (**g**) Total retinal RNA was isolated from mice treated with NPs (n = 5) or PBS (n = 5) at day 14 post-injection. Purified RNA was subjected to RNA-seq and principal component analysis. PC1 = principal component 1; PC2 = principal component 2. (**h**) Heatmap showing gene expression patterns (n = 5) across relevant pathways denoted by the rainbow scale: Red = photoreceptor and RGC health; yellow = oxidative stress response; green = apoptosis; light-blue = chemokines and leukocyte recruitment; purple = microglia activation; dark-blue = glia reactivity. Gene names are listed below the heatmap. The color bar on the right shows the fold-change in gene expression.

Furthermore, we evaluated the expression and distribution of the glial reactive markers GFAP and Iba1 in mouse eyes (Supplementary Fig. S17). Immunofluorescence analysis of retinal sections revealed no appreciable differences in GFAP or Iba1 staining intensity or pattern, suggesting that the NPs do not elicit a detectable inflammatory response in the retina.

To further assess tolerability and safety, we performed RNA-seq on retinal samples from eyes injected with either NPs or PBS. Principal component analysis demonstrated overlapping clustering of the two groups, with no statistically significant separation (Fig. 6g). Likewise, analysis of gene expression patterns across relevant pathways, including photoreceptor and RGC health, cell death, microglia activation and glia reactivity, revealed no statistically significant differences between eyes treated with NPs or PBS (Fig. 6h). Consistent with the absence of changes in GFAP and Iba1 staining, differential expression analysis showed no significant differences in *Gfap* or *Aif1* expression, respectively, between NP- and PBS-injected eyes (p = 1), indicating that the treatment did not induce glial activation. Notably, no expression of pro-inflammatory cytokines (*Il1b*, *Tnf*, and *Il6*) or the inflammasome component (*Nlrp3*) were detected in any of the groups. The complete list of differentially expressed genes is provided in Supplementary Data S1.

In addition, to further support the ocular safety profile, we investigated how NP exposure affected retinal cell viability. R28 (rat retinal neuron cell line) and ARPE-19 (human adult retinal epithelium cell line) cells were exposed to hg-C_3_N_4_ NPs. Viability was evaluated 48 h later using an MTT assay. In general, the NPs were well tolerated by both R28 and ARPE-19 cells (Supplementary Fig. S18a). IC₅₀ values could not be determined, as the dose-response curves did not reach 50% inhibition within the tested concentration range (0–150 µg/cm^2^). Across all time points, cell viability remained above approximately 70%, indicating that the treatment induced only mild cytotoxic effects under these conditions. At 48 hours of exposure, the approximated IC₁₀ was 9.5 and 43.8 µg/cm^2^, for the ARPE-19 and R28 cells, respectively. Similarly, LDH assay (Supplementary Fig. S18b) also confirmed no cytotoxicity by the hg-C_3_N_4_ NPs treatment at a standard to high concentration of 2.0 µg/cm^2^ with 10 minutes of light treatment (10 mW/cm^2^, 1 Hz, 100 ms pulse width). These findings provide additional evidence of substantial tolerability of biosynthesized NPs by ARPE-19 and R28 retinal cells. Moreover, light-induced biomodulation at the cellular level was confirmed on retinal cells by calcium imaging of ARPE-19 cells treated with nanoparticles (Supplementary Fig. S19 and Supplementary Video S28). Collectively, our findings provide strong evidence that the NPs exhibit no cytotoxic or pro-inflammatory activity in retinal cells and tissue, demonstrating that they are both safe and well tolerated.

Next, to evaluate whether the intervention could restore light sensitivity in visually impaired mice (Fig. 7a-h), we administered the NPs into *Pde6β*^rd10^ mice intravitreally. These mice (hereafter rd10 mice) represent a translationally relevant mouse model of autosomal recessive retinitis pigmentosa and typically lose electroretinogram (ERG) responses, a standard measure of photoreceptor function, by ∼8 weeks of age ^69^. Both eyes of 12-week-old rd10 mice were injected with hg-C_3_N_4_ nanoparticles (n = 7) or control particles (SiO₂, n = 3; Red F, n = 4), with OCT and fundoscopic monitoring. Age-matched C57BL/6JRj WT mice (n = 6) served as healthy controls.

**Fig. 7.**
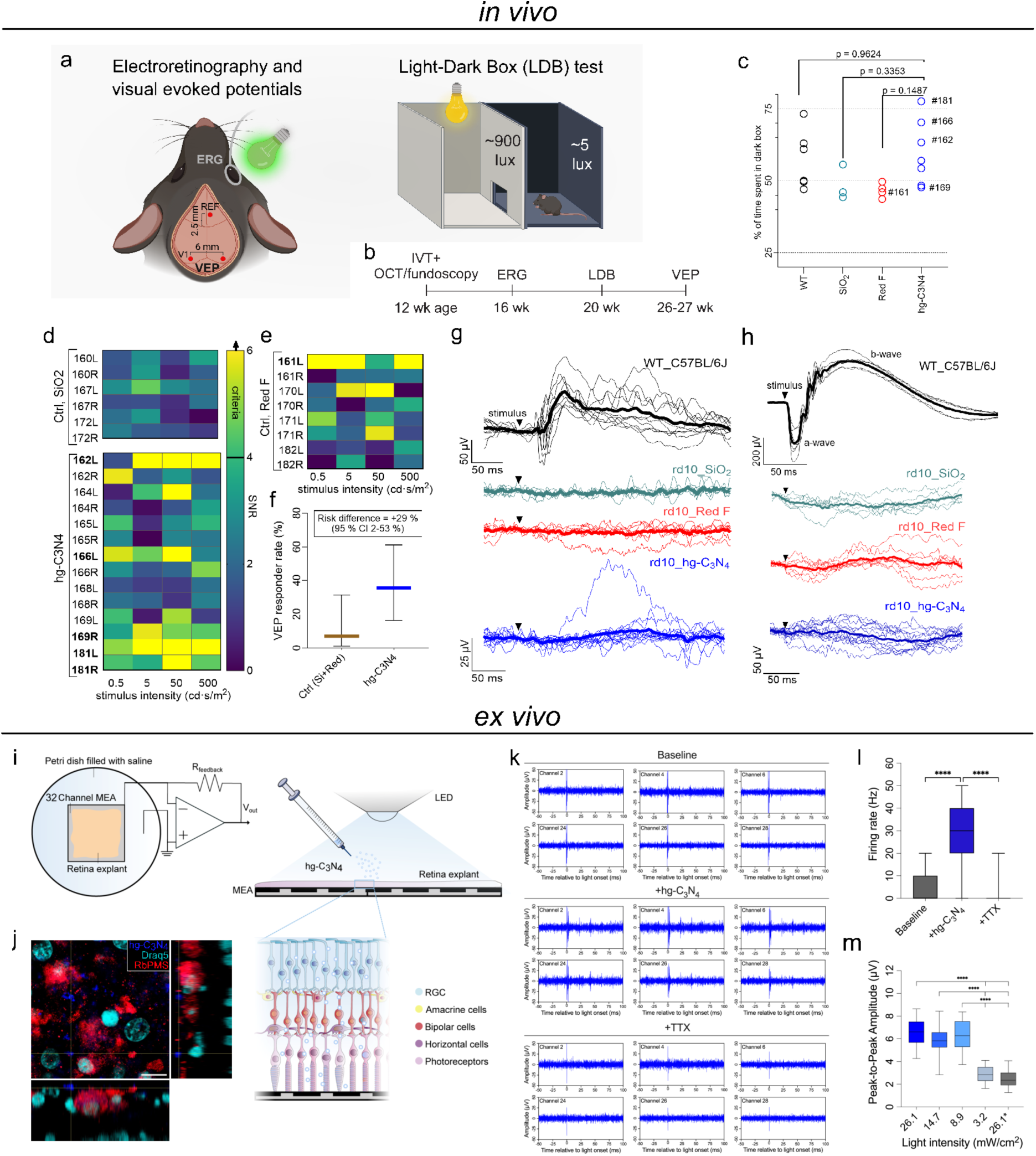
hg-C_3_N_4_ NPs can mediate light responses in vivo and ex vivo. (**a**) Schemes of visual function experiments in rd10 and WT mice *in vivo*. Illustrations created with BioRender.com (**b**) The schedule of experiments in rd10 mice. (**c**) Analysis of dark preference in the light-dark box (LDB) test. Statistical analysis was performed with one-way ANOVA and Tukey’s test for multiple comparisons. (**d-e**) Signal-to-noise ratio (SNR) analysis of VEP responses in rd10 mice. Eyes that reached the criteria as being responders are bolded (criteria: SNR positive ≥ 4.0 in ≥ 3 of 4 stimulus intensities, positive energy ratio ≥ 0.6, peak latency between 80–220 ms, template r ≥ 0.4). (**f**) Fraction of eyes classified as VEP responders. Bars show mean ± Wilson 95 % confidence interval (CI). hg-C_3_N_4_ treatment restored VEPs in 36 % of eyes vs 7 % in controls (risk difference +29 %, 95 % CI 2–53 %). (**g**) Visual evoked potentials (VEPs). Thick lines represent group-mean VEP waveforms whereas the thin lines represent responses from individual eyes. Note different y-axis scale for WT and rd10 traces. (WT, n=10 eyes, N=6 mice; rd10_SiO^2^, n=6 eyes, N=3 mice; rd10_Red F, n=8 eyes, N=4 mice; rd10_hg-C_3_N_4_, n=14 eyes, N=7 mice). (**h**) Representative scotopic ERG waveforms from C57BL/6J WT and rd10 mice in response to a rod-saturating stimulus. WT mice show robust ERG responses, whereas rd10 mice display flat responses across all treatment groups. Note different y-axis scale for WT and rd10 traces. llustrations were created with BioRender.com. (**i**) Schematic of the ex vivo MEA retina recording setup. Inset illustrates RGCs interacting with hg-C_3_N_4_ NPs. (**j**) Fluorescent image of porcine retinal flat mounts showing hg-C_3_N_4_ particle localization on the RGC layer. Scale bar is 10 μm. Blue = hg-C_3_N_4_; turquoise = Draq5; red = RbPMS. (**k**) Representative MEA traces (−50 to +100 ms around light onset) recorded before particle application, after hg-C_3_N_4_ addition, and after TTX. (**l**) Firing frequency 100 ms following light stimulation. Statistical analysis by Kruskal-Wallis test with Dunn’s multiple comparisons with p < 0.0001 denoted as ****. (**m**) Peak-to-peak amplitudes from MEA light dose-response recordings (100 ms light pulses). Statistical analysis by one-way ANOVA with Tukey’s test with p < 0.0001 denoted as ****.

To assess whether hg-C_3_N_4_ treatment could restore visually guided behavior, we tested the mice in a light–dark box (LDB) paradigm four weeks after intravitreal injection (Fig. 7a-b). In this assay, animals freely explore two connected compartments that are identical except that one is brightly illuminated and the other is dark. Mice that perceive light typically prefer the dark compartment, whereas visually impaired mice spend similar amounts of time in both. Among the hg-C_3_N_4_-injected rd10 mice, 3 of 7 spent more than 60% of the time in the dark compartment, a preference level comparable to that observed in 3 of 6 WT animals. In contrast, control rd10 mice injected with SiO₂ or Red F nanoparticles displayed no clear dark preference, consistent with a complete loss of light sensitivity (Fig. 7c).

To independently evaluate light responses in the visual cortex, we next recorded visual evoked potentials (VEPs) using skull screw electrodes placed over the primary visual cortices under urethane anesthesia (Fig. 7d-g). rd10 mice injected with control nanoparticles (SiO₂ or Red F) generally lacked discernible VEPs; however, one Red F-injected eye met the predefined responder criteria (Fig. 7d-e). In contrast, 5 of 14 eyes from hg-C_3_N_4_-treated rd10 mice exhibited reproducible VEP waveforms that met the objective responder definition (SNR ≥ 4, positive polarity, 80-220 ms latency, and ≥ 3 of 4 stimulus intensities passing). Some responses in hg-C_3_N_4_-treated eyes were clearly distinguishable from noise (Supplementary Fig. S20), albeit much smaller and slower than those in WT mice.

Quantitatively, the fraction of VEP-responding eyes in the treated group was 36 % (95 % CI 17–59 %), compared with 7 % (95 % CI 1–31 %) in the combined control group, as estimated using Wilson binomial confidence intervals (Fig. 7f). The risk difference between groups was +29 % (95 % CI 2–53 %), indicating that hg-C_3_N_4_ treatment increased the probability of a measurable cortical light response by roughly one-third. These electrophysiological findings, together with the independently assessed LDB behavior, indicate that a subset of treated rd10 mice recovered both behavioral and cortical sensitivity to light. Notably, the three hg-C_3_N_4_-treated mice that showed dark preference in the LDB test also exhibited the highest VEP SNRs and were classified as electrophysiological responders, suggesting a convergent relationship between behavioral and cortical recovery.

To determine whether the behavioral and cortical light responses observed following hg-C_3_N_4_ treatment were mediated by restored photoreceptor function, we performed scotopic ERG recordings one month after injections under ketamine and medetomidine (KM) anesthesia. No detectable ERG signals were observed in any of the injected rd10 mice, suggesting a lack of light sensitivity from the photoreceptors (Fig. 7h). Of note, the flash ERG response primarily arises from photoreceptor hyperpolarization (a-wave) followed by subsequent bipolar cell depolarization (b-wave); in contrast, RGC electrical activity is not detected due to similar amounts of ON and OFF pathway activation leading to signal cancellation ^70^. Thus, the absence of ERG responses is consistent with the behavioral and cortical light responses arising downstream of classical photoreceptor-driven retinal signaling.

To explore potential retinal factors that might contribute to the observed variability, we examined the residual photoreceptor population. Post-mortem histology (Supplementary Fig. S21) and M-opsin staining (Supplementary Fig. S22) revealed small remnants of M-cones in rd10 retinas. In all treatment groups, M-opsin was confined to the cell bodies rather than the outer segments, and both inner and outer segments were absent. Although classical phototransduction requires intact outer segments, recent studies have shown that cones lacking outer segments can retain light responsiveness ^71^. Moreover, studies in rd10 mice indicate that residual cone-driven responses can persist until approximately P140 (20-week-old), whereas by P238 (34-week-old) this activity is lost ^72^. Thus, a minimal degree of cone-mediated sensitivity cannot be completely excluded in our rd10 mice that underwent VEP recording at 26–27 weeks of age. However, because the extent of cone remnant preservation and M-opsin mislocalization was comparable across all treated and control groups (Supplementary Fig. S21–S22), residual cones alone are unlikely to account for the pattern of VEP and LDB outcomes observed across groups. Taken together, these in vivo findings highlight both the potential and the limitations of hg-C_3_N_4_–mediated vision restoration in the rd10 mouse model, motivating evaluation of retinal light sensitivity under conditions where delivery constraints are minimized and retinal access is fully controlled.

To further evaluate hg-C_3_N_4_–mediated light responses under controlled condtions, we examined an ex vivo preparation in which nanoparticle delivery and retinal exposure can be precisely defined. Here, porcine retinal tissue was placed photoreceptor-side down on a multielectrode array (MEA), with RGCs oriented toward the electrodes (Fig. 7i-j). This configuration enabled direct monitoring of RGC spiking activity in response to controlled optical stimulation, independent of the anatomical barriers and intraocular delivery limitations inherent to the mouse eye.

Applying 1 ms visible light LED pulses (450 nm), we first recorded baseline light-evoked potentials, which elicited a low firing rate of 5 Hz across all channels (Fig. 7k-l, Supplementary Fig. S23) accompanied by small mean peak-to-peak amplitudes (mean 11.6 µV; Supplementary Fig. S26a-b). Following application of hg-C_3_N_4_ NPs to the same tissue and their settlement onto the RGC layer (Fig. 7j), illuminating with the same light dose (30 mW/cm^2^) resulted in a pronounced increase in RGC spiking activity, with the mean firing rate increasing from 5 to 29 Hz, determined over the 1-100 ms window following light onset (Fig. 7k-l). In parallel, peak-to-peak amplitudes also increased from 11.6 to 39.0 µV (Supplementary Fig. S26a-b), consistent with enhanced activation of RGCs in contact with hg-C_3_N_4_. A partial contribution from residual photoreceptors cannot be excluded, although photoreceptor responsiveness may be reduced following tissue dissection, transport and preparation.

To confirm that the recorded signals were of biological origin and reflected compound action potentials generated by RGCs, we next applied Tetrodotoxin (TTX) to block voltage gated sodium channels and suppress neuronal firing ^73^. In the presence of TTX, both the firing rate and peak-to-peak amplitude of light-evoked responses were greatly reduced (1.3 Hz and 11.7 µV respectively) (Fig. 7k-l, Supplementary Fig. S25, S26a-b), confirming that the recorded activity indeed arises from action potential firing in RGCs.

Finally, we assessed the light-dose dependence of the MEA responses. In the presence of hg-C_3_N_4_ NPs (+P), light intensities as low as 8.9 mW/cm^2^ were sufficient to produce measureable RGC activity (Fig. 7m). By contrast, peak-to-peak amplitudes recorded at 3.2 mW/cm^2^ with particles were comparable to those obtained at 26.1 mW/cm^2^ in the absence of particles (-P), indicating a substantial shift in light sensitivity. Light evoked peak-to-peak amplitudes in the dose-response experiment were lower overall than those observed in the initial recordings, likely reflecting both reduced illumination levels and variability in the physiological state of the individual porcine retina preparations.

## Discussion

In summary, we have demonstrated the versatility of chloroplast-mimicking hg-C_3_N_4_ NPs for multiscale photo-modulating electrophysiology, at the subcellular level, intercellular level, and tissue level. Tools to probe intracellular and intercellular electrophysiological mechanisms are crucial for studying physiological and pathological pathways, not only prevailing in cardiovascular and neurodegenerative diseases. For instance, a recently discovered hub of highly connected glioma cells, similar to heart ‘pacemaker’ cells, has been shown to drive calcium propagation through tumor microtube networks and promote expansion of the lethal malignant glioma via activation of MAPK and NF-κB signaling pathways ^74^. At present, these studies are often hampered by the available disruptive methods ^75^. We present the hg-C_3_N_4_ as organic semiconductor NPs to leadless modulation of single cell signaling across a broad spectrum of excitable and non-excitable cell types and endow intercellular signal propagation by applying laser stimulation with subcellular resolution. Accumulatively, the capability of hg-C_3_N_4_ NPs to pace cardiomyocytes network’s synchronous beating was achieved by simple LED stimulation. The intercellular propagation of these signals is likely facilitated by gap junction coupling, a hallmark feature of cardiac tissue that ensures coordinated contraction. Though being out of the scope of the current work, application of optical-fiber-coupled endoscope ^76^ should further enable in vivo heart pacing and clinical translation.

The hg-C_3_N_4_ NPs could be safely internalized, an important feature for stimulation on the intracellular level. Combined with the clear directedness quantified as cos(θ) (Fig. 3f), as well as the clustering of particles around the nucleus (Fig. 3e, Supplementary Fig. S4c), the inhibitor assay results indicate that the particles may internalize through multiple pathways, including both phagocytosis- and endocytosis-driven mechanisms (Fig. 3g) ^48,49^, and are actively trafficked in endosomal compartments towards the perinuclear region ^50,77^. Importantly, using an NP dose that caused no changes in the retinal gene expression profile (Fig. 6h) and maintained 99 % viability in R28 cells, we achieved a partial restoration of light responsiveness in rd10 mouse retinas, demonstrating therapeutic benefit without detectable cytotoxicity.

The photo-response mechanism of hg-C_3_N_4_ NPs has been revealed to be of both anodic faradaic and photothermal nature. The mechanism behind the observed intracellular calcium transient is attributed to hg-C_3_N_4_ NPs interacting with calcium storing organelles like the endoplasmic reticulum (ER) (Fig. 1b, 4a) ^78^. As we could show that the photo faradaic and photothermal effect go along with ROS production (Fig. 2, Supplementary Fig. S2), the generated ROS activates intracellular RyRs, (Supplementary Fig. S10) leading to ER calcium release. In excitable cells, the process is independent of IP₃R signaling, extracellular calcium influx, TRP channels, and thermal effects, reinforcing that intracellular ROS-triggered ER calcium release is the central mechanism of stimulation in this system^79,80^.

Compared with silicon nanowires, a widely explored optoelectronic interface ^81^, hg-C_3_N_4_ generate anodic rather than cathodic faradaic photocurrents and exhibit high chemical stability (Supplementary Fig. S1b, S16) ^19^, representing a unique advantage to silicon nanowires which have been shown to gradually degrade under physiological conditions ^81^. Consistent with this, the photocurrent generated by hg-C_3_N_4_-coated substrates remained stable under repeated 473 nm laser pulses across multiple on–off cycles. (Supplementary Video S13). Together, these intrinsic properties support the feasibility of hg-C_3_N_4_ for repeated and long-term use in biological environments.

The anodic currents leading predominately to oxidative reactions and photothermal effects, both contribute to the generation of H_2_O_2 80_. While interfacial reactions are complex and additional pathways for ROS generation may exist ^82^, ROS such as H_2_O_2_ are important signaling molecules involved in many fundamental physiological processes ^83^, including interactions with neuronal ion channels ^84,85^. Although physiological ROS levels are tightly regulated around 1–5 µM and elevated concentrations can be cytotoxic ^83,86^, neither cytotoxic effects (Supplementary Fig. S5, S6, S18) nor inflammation induction *in vivo* (Fig. 6g-h, Supplementary Fig. S17) ^87^ were observed in this study.

Photostimulation is a concept worthy of further development to achieve high spatiotemporal resolution. The demonstrated feasibility of safe and therapeutic *in vivo* delivery to the murine retina positions hg-C_3_N_4_ NPs as a potential injectable retinal prosthesis, with the capability to restore light sensitivity in patients with severe photoreceptor loss due to e.g. inherited retinal degenerations or age-related macular degeneration. However, in rd10 mouse experiments, relatively high luminance levels were required to elicit detectable light responses in the visual cortex. This limited sensitivity may reflect to the limited efficiency of intravitreal delivery in bringing NPs into sufficient contact with RGCs. To further optimize this vision restoration strategy, alternative delivery methods, such as subretinal injection, should be explored. These approaches may facilitate NP integration with bipolar cells, enabling signal amplification and more natural retinal processing compared to direct RGC targeting ^88^.

Finally, although the low tissue penetration depth of blue light might limit the use of hg-C_3_N_4_ NPs in certain applications, red-shifting the absorbance of g-C_3_N_4_ toward near infrared has been proven to be feasible ^89^. Additionally, given that focused ultrasound has been utilized to generate ROS for vinyl monomer polymerization ^90^, the emerging piezoelectric properties of g-C_3_N_4 91_ could further expand the applicability of hg-C_3_N_4_ NPs for deep tissue stimulation.

## Methods

### Synthesis of silica template

The silica template was synthesized according to the classical Stöber method ^27^. In detail, the mixed solution containing 4.0 mL of aqueous ammonia (28 wt%), 74.0 mL of ethanol and 10.0 mL of deionized water were stirred vigorously at 30 °C for 1h. Then, 5.6 mL of tetraethoxysilane (TEOS) was slowly added to the above mixture under continuous stirring and stood still for 1 h to yield uniform monodisperse silica cores. Next, 3.56 mL of TEOS and 1.71 mL of n-octadecyltrimethoxysilane (C_18_TMOS) were added dropwise to the above solution while stirring. After the solution was dropped, the mixture was continued to stir for 15 min to form a thin silica shell around the dense silica core. Subsequently, the mixed solution was allowed to stand for 3 h to promote the cohydrolysis and condensation of the TEOS and C_18_TMOS. The mixed solution was centrifuged, washed in DI water and ethanol, dried at 80 °C and calcined at 550 °C for 6 h in air. Finally, the obtained silica template was neutralized with a 1 M HCl solution and dried at 80 °C overnight.

### Synthesis of hg-C_3_N_4_ NPs

A solution of 1.0 g of the silica template and 5.0 mL of cyanamide aqueous solution (50 wt%) was mixed for 30 min in a flask, then the mixture was ultrasonicated in vacuum at 60 °C for 3 h and stirred at 60 °C overnight. After centrifugation and drying, the products were transferred to chemical vapor deposition (CVD) furnace for calcination and heated to 550 °C under flowing N_2_ for 4 h with a heating rate of 4.4 °C min^−1^. The obtained products were treated with 4 M NH_4_HF_2_ for 12 h to remove the silica template. In the last, the yellow powders were centrifuged, washed three times in DI water and once in ethanol, and then dried in vacuum at 60 °C for 10 h to obtain the final HCNS products.

### Electron microscopy

The particles morphology was analyzed by SEM (TM3030Plus, Tabletop microscope, HITACHI). Transmission electron microscopy (TEM, Tecnai G2 Spirit) was used to visualize the hollow-sphere structure. TEM copper grids (Merck) were employed for analysis with an acceleration voltage of 120 kV. Transmission electron microscopy with energy-dispersive X-ray spectroscopy (TEM-EDS) was performed using JED-2300 F200 microscope operated at an accelerating voltage of 200 kV. Samples were dispersed in ethanol, then dropped-cast onto 300-mesh copper grids coated with amorphous carbon film dried at room temperature prior to measurements.

### Biodegradation characterization

The sample was dispersed into PBS buffers (pH 4.5 and 7.0) and drop-cast onto the conductive tape at days of 0, 7, 14 and 30. After drying at room temperature, the samples were sputter-coated with 10-nm Au layer for SEM characterization. We acknowledge iMAT center at Aarhus university for access to the Clara SEM facility.

### Dynamic light scattering

The hydrodynamic size distribution of nanoparticles was measured by dynamic light scattering (DLS) using Malvern Zetasizer Nano ZS. Samples were dispersed in filtered PBS buffer (pH 7.4), diluted to 0.05 mg/mL in filtered PBS buffer and DMEM with 10% FBS and equilibrated at 25 °C for 3 min before measurement. The intensity-weighted size distribution was derived from the average of 3-5 runs. Zeta potential was measured using the same instrument for 5-8 runs.

### UV-vis spectroscopy

UV-vis spectra were acquired with the GENESYS 150 UV-Visible Spectrophotometer (Thermo Fisher Scientific).

### X-ray diffraction

X-ray diffraction (XRD) measurements were performed using SmartLab 9 kw X-ray diffractometer (Cu K ɑ radiation) operated at 60 kV and 220 mA. The data were collected in the 2θ range of 5°-60° with a scanning step of 0.02° and a scan speed of 20° min^-1^. The samples were ground into fine powders and pressed onto a glass slide for measurement.

### X-ray photoelectron spectroscopy

X-ray photoelectron spectroscopy (XPS) measurements were carried out on Thermo Scientific ESCALAB Xi+ XPS spectrometer with Al K ɑ radiations, and with the C 1s peak at 284.8 eV as an internal standard for all the spectra. The XPS spectra peak deconvolution was performed with Avantage software using a Shirley background.

### Photothermal Response Measurement

Photothermal effects were quantified following a previously established protocol, using a standard patch-clamp setup equipped with an upright microscope (BX61WI, Olympus) and a 20×/0.5 NA water-immersion objective for optical excitation. Illumination was provided by a 625 nm LED (M625L4, Thorlabs) with a ∼750 µm spot size. Light pulses were TTL-triggered via a Digidata 1550 digitizer (Molecular Devices). Glass micropipettes (∼1 MΩ, pulled with a P-97 puller, Sutter Instrument), filled with 1× PBS, were positioned approximately 2 µm above the photoactive material immersed in the same PBS solution. Ionic currents were recorded in voltage-clamp mode using an Axopatch 200B amplifier and pClamp software. The photothermal response was determined by plotting the light-induced current change (ΔI_light) against the pipette holding potential (I0) and extracting the slope. Local temperature increase was estimated by calibrating pipette resistance changes in pre-heated PBS solutions (20–50 °C), using a thermocouple placed near the pipette tip to establish a resistance–temperature calibration curve.

### Electron paramagnetic (spin) resonance (EPR/ESR)

Generation of reactive oxygen species (ROS) was assessed using electron spin resonance spectroscopy (Bruker Magnettech ESR5000). Spin traps were used to detect specific species: TEMP (100 mM in H₂O) for singlet oxygen (^1^O₂), DMPO (100 mM in methanol) for superoxide (•O₂^−^), and DMPO (100 mM in water) for hydroxyl radicals (•OH). Nanoparticles (hg-C_3_N_4_) were dispersed at 1 mg/mL in either DI water or methanol using probe sonication. Each dispersion was mixed 1:1 with the relevant spin trap and loaded into sealed glass capillaries. Samples were illuminated using an LED light source (450 nm, 75 mW/cm²) for 0, 5, and 10 min. ESR spectra were recorded immediately after using the following parameters: magnetic field range (B) = 330.0–340.0 mT, center field (B₀) = 335.0 mT, sweep width = 10 mT, sweep time = 30 s, modulation = 0.2 mT, and modulation frequency = 100 kHz.

### Cell line culture

NIH/3T3 mouse fibroblasts (ATCC CRL-1658, Manassas, VA, USA), R28 rat retinal precursor cells (Kerafast, Newark, CA, USA), ARPE-19 human retinal pigment epithelial cells (ATCC CRL- 2302), and HeLa human cervical cancer cells (ATCC CCL2, Manassas, VA, USA) were cultured in Dulbecco’s Modified Eagle Medium (DMEM) supplemented with 10% fetal bovine serum (FBS), 1% P/S, and 4 mM L-glutamine. All cells were kept at 37 °C and 5% CO_2_ and media were changed every second day.

HL-1 murine cardiac muscle cells (Merck SCC065) were cultured according to the manufacturer’s protocol. Before seeding, the flasks and wells were pre-coated with a Gelatin/Fibronectin solution (EMD Millipore ES-006, Sigma F-1141) for at least 2 h. HL-1 cells were cultured in Claycomb Basal Medium (Sigma, 51800C) supplemented with 10% HL-1 qualified FBS (EMD Millipore, TMS-016-B), 2 mM Glutamax (ThermoFisher), 0.1 mM norepinephrine (Sigma, A0937), and 1% P/S. The cells were kept at 37 °C in a humidified 5% CO2 atmosphere, and the culture medium was changed daily.

### Cardiac cell culture

Primary cardiomyocytes (CMs) and cardiac fibroblasts (CFs) were isolated and harvested from P0-5 neonatal rats following literature ^78^ and reagents were obtained from Pierce™ primary cardiomyocyte isolation kit (Thermo Fisher Scientific) ^92^. In short, the heart tissues were excised into ice cold Hanks’ Balanced Salt solution (HBSS) medium without Ca^2+^ or Mg^2+^ and digested with reconstituted papain and thermolysin enzymes. Isolated CFs/CMs hybrid cells were washed with HBSS, seeded onto fibronectin (Sigma)-coated glass bottom imaging dishes, and cultured in high glucose DMEM supplemented with 10% FBS, 1% penicillin–streptomycin, and 1% GlutaMAX. Animal procedures were approved by the University of Chicago Institutional Animal Care and Use Committee (IACUC) and conducted in complete compliance with the IACUC Animal Care and Use Protocol.

### iPSC-CM culture

Human induced pluripotent stem cell–derived cardiomyocytes (iPSC-CMs) were kindly provided from Prof Manuel Maria Mazo Vega from Cima Universidad de Navarra. Cells were seeded at a density of 250,000 cells/cm² in seeding medium consisting of RPMI 1640 (Sigma, R8758), 10% KnockOut™ Serum Replacement (KSR; Gibco, 10828010), 2% B27 supplement (Thermo Fisher Scientific, 17504044), 1% penicillin-streptomycin (P/S), and 10 µM Y-27632 ROCK inhibitor. Cells were plated in 24-well plates pre-coated with Matrigel (Corning, 354230). The following day, the medium would be changed to maintenance medium composed of RPMI 1640, 2% B27, and 1% P/S. Cells were cultured at 37 °C in a humidified 5% CO₂ incubator, and the medium was refreshed every 2–3 days.

### Particle co-culture

Prior to *in vitro* experiments, hg-C_3_N_4_ NPs were dispersed in PBS at 1 mg mL⁻¹ by probe sonication (MS73, 60 % amplitude, 1 s on/off, 3 × 5 min). For sterilization, NPs were either autoclaved, sonicated in ethanol followed by washing and resuspension in PBS, or exposed to UV light for 15 min. Unless stated otherwise, hg-C_3_N_4_ NPs were added to cell cultures at a surface dose of approximately 2.0 µg/cm^2^ hg-C_3_N_4_.

### DCFH-DA assay

To assess intracellular ROS production, a DCFH-DA assay (Abcam, ab113851) was performed following the manufacturer’s protocol. NIH/3T3 cells were seeded in 96-well plates at a density of 20,000 cells/well and incubated overnight to allow cell attachment. hg-C_3_N_4_ nanoparticles were added 8 hours prior to the assay at a concentration of 2.0 µg/cm². Cells were then subjected to continuous blue light stimulation (450 nm, 10 mW/cm²) for 10 minutes using an LED array.

Immediately following stimulation, 20 µM DCFH-DA working solution was added to each well, and the cells were incubated for 45 minutes at 37 °C in the dark. After incubation, the dye solution was aspirated, and the wells were washed twice with warm PBS. Fluorescence was first assessed qualitatively using a green fluorescence filter on the EVOS M7000 microscope. Quantitative measurements were then performed 4 hours post-staining using a microplate reader (excitation: 485 nm, emission: 535 nm). Four experimental groups were evaluated (w/o NPs, w/o light), each in quadruplicates.

### LDH cytotoxicity assay

Cytotoxicity was evaluated using a lactate dehydrogenase (LDH) assay kit (Roche) following the manufacturer’s instructions. NIH/3T3 cells were seeded at 2,500 cells/well, ARPE-19 and R28 cells at 5,000 cells/well, and iPSC-derived cardiomyocytes (iPSC-CMs) at 80,000 cells/well, all in 96-well plates and in sextuplicates. For NIH/3T3 cells, two experimental groups were included: with and without nanoparticles (NPs). For ARPE-19, R28, and iPSC-CMs, two additional groups were added: with and without light exposure. Where applicable, hg-C_3_N_4_ nanoparticles were added at a concentration of 2.0 µg/cm². Light stimulation was performed using a blue LED array (10 mW/cm², 1 Hz, 100 ms pulse width) for 10 minutes, administered 2 hours before the assay.

After 24 hours of incubation, 80 µL of the cell culture medium was collected and centrifuged at 1700 rpm for 5 minutes at 4°C. The activity levels of lactate dehydrogenase (LDH) were then measured by transferring 50 µL of the supernatant to a new 96-well plate and adding reaction mix containing catalyst and dye in a 1:1 ratio. Subsequently, the plate was incubated in the dark at room temperature for 30 minutes, followed by analysis of absorbance on a Victor Multilabel Reader (PerkinElmer, US) at 490 nm. To establish the toxicity levels, the LDH activity of cells cultured without particles was regarded as the low toxicity control, while that of cells treated with 1% Triton X-100 was marked as high-toxicity control. The relative toxicity levels were calculated using the below equation:

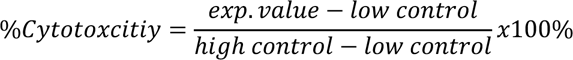

### Live/Dead staining

Cell viability upon exposure to particles and light was assessed using Live/Dead staining using NIH/3T3 cells cultured for 24 h. The particles (2.0 µg/cm^2^) were introduced 6 hours prior to the light stimulation. The staining solution, a combination of 1:1000 Calcein-AM (green dye) and 1:500 Ethidiom iodide (red dye) medium, was added after aspiration of the culture medium and a single rinse with serum-free medium. After adding the staining solution, cells were then incubated for 30 min at 37°C. Afterwards, the staining solution was discarded, wells were rinsed with PBS, and cells could be examined using a fluorescent microscope (Invitrogen EVOS M7000).

To evaluate long-term cytocompatibility under repeated stimulation, Live/Dead staining was additionally performed on ARPE-19 cells, R28 cells, and iPSC-derived cardiomyocytes after 7 days of culture with bidaily blue light stimulation (10 mW/cm², 1 Hz, 100 ms pulse width, 10 minutes).

### CCK-8 proliferation assay

The effect of particles on cell proliferation was assessed using a Cell Counting Kit-8 (CCK-8, Dojindo). 5000 NIH/3T3 cells were seeded in 96-well plates in sextuplicate. 2.0 µg/cm^2^ of hg-C_3_N_4_ particles were added. The CCK-8 assay was performed on days 1, 3, and 8 following the manufacturer’s protocol. Briefly, the culture medium in each well was replaced with CCK-8 solution diluted 1:20 in medium. The cells were then incubated for 2 hours, after which the medium was transferred to a new 96-well plate and fresh culture medium was replenished. Subsequently, the absorbance at 450 nm was analyzed on Victor Multilabel Reader (PerkinElmer, US). The proliferation index is calculated as:

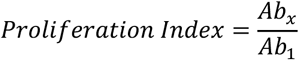

with Ab being the absorbance value of the CCK-8 assay measured on day x (x = 1, 3, 8).

### MTT assay

ARPE-19 cells were seeded in a transparent 96-well plate, grown to confluency and serum-starved. R28 cells were serum-starved and seeded at a density of 25.000 cells/well in a transparent 96-well plate. The next day, the cells were incubated with 5, 25, 50, 100, or 150 µg/cm^2^ of hg-C_3_N_4_ NPs. Cells treated for 1 h with 4% Triton were the reference of 100% toxicity. Untreated cells (no NPs) served as the 100% reference for percent-of-control calculations. 48 h after treatment with hg-C_3_N_4_ NPs, the MTT assay was conducted using the MTT cell proliferation assay kit (Cayman Chemical, Ann Arbor, MI, USA) following the manufacturer’s instructions. Absorbance was measured at 570 nm using the SpectraMax iD3 absorbance reader (Molecular Devices, San Jose, CA, United States). IC₁₀ values were obtained directly from the curves by interpolating the concentration at which cell viability reached 90%. Each treatment condition was tested in quintuplicates, and viability values represent the mean (±SD) of individual measurements relative to control (no NPs).

### hg-C_3_N_4_ NP internalization

For live cell imaging and single particle tracking (SPT) of NIH/3T3 and R28 cells an inverted spinning disk confocal microscope (SDCM (Olympus SpinSR10, Olympus, Tokyo, Japan) was applied. The SDCM used an oil immersion 60x objective (Olympus) for recording images and videos with a total pixel width of 183 nm and numerical aperture of 1.4, and a CMOS camera (photometrics PRIME 95B). Excitation wavelength of 640 nm was used to excite the CellMask DeepRed plasma membrane stain (Invitrogen, C10046) at 2% laser power, 100 ms exposure time while hg-C_3_N_4_ NPs where excited with a 488 nm laser at laser power at 25%, 100 ms exposure time for 100 frames (frame rate 102 ms) with Gain 2.

7.000 NIH/3T3 or 10.000 R28 cells per well were added with the appropriate amount of growth media and left for incubation at 37°C, 5% CO_2_ the day prior to imaging on IbiTreat microscope plates. On the day of microscopy, 100 µL growth media was replaced with 100 µL test solution of hg-C_3_N_4_ NP dispersion for a final concentration of 0.6 µg/cm^2^ and incubated for the given amount of time at 37°C and 5% CO2. The test solution was then removed and replaced with pre-heated 200 µL 1X CellMask DeepRed plasma membrane stain and incubated for 5 min, before the wells were washed 1x with imaging media and following replaced by fresh imaging media for data acquisition. The inhibition study was performed similarly, but before addition of the of hg-C_3_N_4_ NP test solution, the cells were incubated with the given inhibitors for 1 hour (500 uM Amiloride (ThermoScientific^TM^, 15414599), 1 uM Bafilomycin (MedChemExpress, HY-100558), 100 uM Chloroquine (Sigma-Aldrich, C6628), 80 uM Dynasore (Sigma-Aldrich, D7693), 10 ng/mL Nystatin (Gibco^TM^, 11548886) or DMEM+10% FBS for the control). Subsequently, 0.6 µg/cm^2^ hg-C_3_N_4_ NP dispersion was added for 3 hours in the presence of the inhibitors. Finally, the test solution was removed and replaced with pre-heated 200 µL 1X CellMask DeepRed plasma membrane stain and incubated for 5 min, before replacing with imaging media for data acquisition. For detection and SPT, the Trackpy^93^ package in python based on the Crocker-Grier algorithm^94^ with in house modification from the Hatzakis lab was used with an initial SNR threshold between 0.3–1.0 based on manual inspection for each FOW and a postprocessing local SNR threshold of 0.3 kept constant for all conditions. Cells were segmented using CellPose^95^ and followingly particles were assigned to the cell masks to quantify particle internalization per cell. For investigating the directedness, a 0.5 mg/mL hg-C_3_N_4_ NP stock was prepared in PBS by sonication for 15 min. The stock dispersion was diluted in media and added to the cells to achieve a final particle concentration of 0.6 µg/cm^2^. The internalization videos were recorded using the Nikon TI2-E inverted microscope and NIH/3T3 fibroblasts. Particle tracking of internalized particles was done using Manual Tracking plugin in ImageJ. 3D imaging of internalized hg-C_3_N_4_ NP colocalized with lysosomes in NIH/3T3 cells was imaged on a Zeiss Lattice Light Sheet 7. 0.6 µg/cm^2^ hg-C_3_N_4_ NP was incubated with the cells for 21 hours, after which LysoTracker DeepRed (1:20.000 dilution) was added to the cells and incubated 3 h. Dual imaging of LysoTracker DeepRed (excited with 640 nm laser, laser power 1%) and hg-C_3_N_4_ NPs (excited with a 488 nm laser at laser power at 5%), was performed with a 30x1000 lightsheet and 50 ms exposure time in 539 slices (107.6 µm). The image was deskewed, deconvolved and displayed in Zeiss’ Zen software.

### Calcium imaging and laser stimulation

To visualize intracellular calcium Cal-520, AM (ab171868, abcam) at 4 µM was used. After incubating the cells with dye in glass bottom dishes at 37 °C for 30 min, the cells were rinsed and incubated for another 30 min at 37 °C in medium. The cells were observed with a 60x oil immersion objective using a Nikon TI2-E inverted microscope. Laser (473 nm) power was calibrated using the PM100D optical power meter (ThorLabs) and S120C photodiode power sensor (ThorLabs).

Isochronal and vector maps were generated using the ImageJ plugin Spiky ^96^. An online available macro was used to generate ΔF/F_0_ movies ^97^. Calcium flux videos were quantified with ImageJ by defining regions of interest around cells and measuring mean gray values (F). As background value (F_0_) the mean gray value of an ROI without cells was defined. The mean fluorescence intensity normalized to the background was then calculated as ΔF/F_0_ for each individual frame.

### Pharmacological experiments

To investigate the molecular pathways underlying calcium signaling, we conducted a series of pharmacological blocking experiments in neonatal rat cardiomyocytes co-cultured with hg-C_3_N_4_. Cells were first loaded with 4 µM for 30 min, followed by initial imaging in standard extracellular buffer as a baseline control. Subsequently, cells were treated with specific inhibitors or antioxidants as indicated below, and after a brief recovery period, laser stimulation (473 nm, 100 ms pulse) was applied to non-previously imaged cells.

The following pharmacological agents were employed to dissect the signaling pathway:

- N-acetylcysteine (NAC) (500 µM, 1 h pretreatment) was used to scavenge intracellular ROS.
- Ryanodine (25 µM, 10 min) was applied to inhibit ER ryanodine receptors (RyRs).
- 2-APB (200 µM, 10 min) served as an inhibitor of IP3 Rs receptors.
- Ruthenium Red (RR) (10 µM, 10 min) was used to block thermosensitive TRP channels.

All inhibitors were administered in culture medium. Control groups were imaged in parallel under identical conditions without drug treatment.

For experiments in calcium-free conditions, extracellular buffer was replaced with calcium-free HBSS immediately prior to stimulation. The fluorescence traces presented represent typical single-cell responses across at least 4 independent cells per condition.

### HL-1 pacing

HL-1 cells were cultured in 24-well plates at a seeding density of 100,000 cells per well. Upon reaching 90% confluency, particles were added at concentrations of 2.0 µg/cm^2^ followed by a 6-hour settling period before initiating calcium imaging.

For calcium flux imaging, cells were stained with Fluo-4 AM (Invitrogen, F14201). Samples were rinsed with Hanks’ Buffer (Sigma H6648) supplemented with 0.1 mM norepinephrine (Sigma, A0937), 5 mM glutamax, and 20 mM Hepes (Sigma H0887). Subsequently, cells were incubated with 5 µM Fluo-4 AM diluted in the same buffer for 30 min at 37°C, 5% CO_2_. After rinsing twice, the cells were incubated in the buffer for an additional 20 minutes. Following the aspiration of the buffer, supplemented Claycomb Medium was added and used during imaging.

Stimulation was performed using light irradiation at a power of 10 mW/cm^2^, frequency of 1 Hz, and pulse width of 100 ms. Calcium flux was monitored and recorded via a 10x objective on the EVOS microscope (EVOS m7000) equipped with an onstage incubator (37°C, 5% CO_2_). The fluorescence traces presented represent typical cell responses across at least 6 independent cells per condition.

### iPSC-CM pacing

iPSC-derived cardiomyocytes were cultured in 24-well plates pre-coated with Matrigel at a density of 250,000 cells/cm². Upon forming a confluent monolayer, particles were added at a concentration of 2.0 µg/cm², followed by an 8 h settling period prior to calcium imaging.

For calcium flux imaging, cells were stained with Fluo-4 AM (Invitrogen, F14201). After rinsing, cells were incubated for 30 minutes at 37 °C in Thyrode’s solution (in mM): NaCl 138, KCl 4, CaCl₂ 1.8, MgCl₂ 1, NaH₂PO₄ 0.33, HEPES 10, glucose 10; pH 7.4, containing 5 µM Fluo-4 AM. Cells were then rinsed twice and incubated in fresh Thyrode’s solution for an additional 20 minutes.

Light stimulation was applied at 30 mW/cm², 1 Hz, and 50 ms pulse width. Imaging was performed using an EVOS m7000 microscope with a 10x objective and onstage incubator (37 °C, 5% CO₂). All conditions were performed in triplicate (three independent wells per condition), and fluorescence traces represent typical single-cell responses from each replicate.

### Pharmacological blockers

To investigate the mechanism of light-induced calcium wave propagation during synchronized stimulation of cardiomyocytes, HL-1 cells were cultured with hg-C_3_N_4_ and stained with Fluo-4 AM as described above. Two pharmacological blockers were tested individually: apyrase (50 U/mL) and carbenoxolone (CBX, 200 µM). Blockers were dissolved in a modified HBSS buffer containing (in mM): 1.3 CaCl₂, 0.5 MgCl₂, 0.4 MgSO₄, 5.3 KCl, 0.4 KH₂PO₄, 4.2 NaHCO₃, 137.9 NaCl, 0.3 Na₂HPO₄, 5.56 D-glucose, and 10 HEPES (adjusted to pH 7.4). Blockers were added to the cells 15 minutes prior to light stimulation.

Light stimulation was performed for 10 minutes using the same settings described above. Calcium dynamics were recorded before and during blocker treatment using a 10× objective on the EVOS M7000 microscope, equipped with an on-stage incubator (37 °C, 5% CO₂). Fluorescence traces represent typical single-cell responses, with data collected from at least six independent cells per condition.

### Calcium imaging and LED stimulation of ARPE-19 cells

To evaluate the ability of hg-C_3_N_4_ nanoparticles to induce calcium signaling in retinal cells, human retinal pigment epithelium cells (ARPE-19) were used. Cells were seeded in 24-well plates at a density of 150,000 cells/cm² and incubated overnight. The following day, hg-C_3_N_4_ nanoparticles were added at a concentration of 2.0 µg/cm². Approximately 8 hours after nanoparticle addition, cells were stained with 50 µM Fluo-4 AM in modified HBSS buffer (in mM): 2 CaCl₂, 1.2 MgCl₂, 5 KCl, 0.44 KH₂PO₄, 4.2 NaHCO₃, 137 NaCl, 5 D-glucose, 20 HEPES, with pH adjusted to 7.4. Cells were incubated with the dye for 40 minutes at 37 °C, followed by a wash and a 15-minute quenching step in dye-free buffer.

Calcium transients were recorded using a live-cell fluorescence microscope (EVOS M7000), before, during, and after blue light stimulation (10 mW/cm², 3 s) using an external LED source. Experiments were performed in triplicate across four conditions: w/o NPs and w/o light stimulation.

### Particle tracking

The ONI Nanoimager was employed for single particle tracking in cell free media to investigate stokes diameter and diffusion coefficient. The maximal frame gab was set to 5, maximal distance between frames was 0.7 µm and the minimum number of steps was set to 20. For particle tracking of internalized particles in 3T3 cells the Manual Tracking plugin for ImageJ was used. Based on the tracking data acquired with Manual Tracking plugin, the directedness was calculated as a parameter for directional movement. The angle θ was defined as the angle between particle movement direction and direction of the center of an individual cell. The directedness is defined as the cosine of this angle. If particle movement is directed exactly towards the center of the cell the value is 1, whereas the value 0 indicates perpendicular movement, and -1 movement in the opposite direction. The average directedness was calculated as Σcos(θ)_i_/n, where θ was the angle between the individual movement directionality vector and the directionality vector towards the center of the cell relative to the particle position (n = 8) ^44^.

### Photocurrent measurement

A detailed protocol on how to acquire photocurrent data has been described previously ^32^. Briefly, the setup was based on an electrophysiology patch-clamp aperture. A P-97 micropipette puller was used to pull glass pipettes and achieve a resistance of 2–4 MΩ. A micromanipulator was used to bring a silver chloride electrode inserted in the glass pipette in close proximity to the particles (<10 µm). The sample as well as the objective is immersed in PBS. The upright microscope (Olympus, BX61WI) allows to illuminate the sample with a 365 nm LED light, with a pulse duration of 100 ms. An AxoPatch 200B amplifier (Molecular devices), and Digidata 1550 digitizer (Molecular devices), and Clampex software (Molecular devices), worked together to conduct the measurement and control the light source.

### H_2_O_2_ quantification

Hydrogen Peroxide (H_2_O_2_) production was evaluated using different concentrations of particles in PBS (0, 1, 2, 5 µg/cm^2^). The solutions were added to 24-wells and particles were allowed to settle for 2 hours before being irradiated with 450 nm LED light (75 mW/cm^2^), pulsed at 100 ms duration and 1 Hz frequency for 30 minutes. Control groups were kept light-protected under the same conditions. Subsequently, 3x100 µL were transferred from each well to a black 96-well plate. The aliquots were stained with a solution containing 91 µL of horseradish peroxidase (HRP; 4250 units/L, Thermofischer), p-hydroxyphenyl acid (pOHPAA; 1.0e-3 M, Sigma), in 0.25 M Tris buffer (Sigma). Standards with known H_2_O_2_ concentrations were treated in the same way. Fluorescence intensity was measured with an excitation of 320 nm and an emission of 405 nm using a microplate reader (CLARIOstar Plus Microplate Reader).

### Experimental animals

The use of laboratory animals was approved by the Danish Animal Inspectorate (authorization no. 2020-15-0201-00745) and by the Finnish Project Authorization Board (ESAVI/26320/2021). For experiments conducted in Aarhus, eight-week-old C57BL/6JRj male mice were purchased from Janvier Labs (Le Genest-Saint-Isle, France). C57BL/6J mice (n=3 females, n=3 males) used at the University of Eastern Finland (UEF) originated from the colony of Lab Animal Centre at UEF. The original colony of the retinitis pigmentosa model B6. CXB1-Pde6brd10/J (RRID: IMSR_JAX:004297, referred to as rd10), mice were a kind gift of Dr. Thierry Leveillard (Sorbonne University) but had been bred as a homozygote line at for at least five generations. In total fourteen rd10 mice were used in this study (n=5 females and n=9 males, randomly divided between groups). Animals were maintained under a 12–/12-hour light/dark with free access to autoclaved tap water and standard chow. All procedures were conducted in accordance with the Directive 86/609/EEC for animal experiments, FELASA Guidelines and Recommendations, and ARVO Statement for the Use of Animals in Ophthalmic and Vision Research.

For injections and eye imaging mice were anesthetized with an intraperitoneal injection of a mixture of ketamine and medetomidine (KM) hydrochloride (Ketador or Ketaminol Vet [60–100 mg/kg;, Richter Pharma AG, Wels, Austria; or MSD Animal Health, Espoo, Finland] and Cepetor or Domitor Vet [0.5–1 mg/kg; ScanVet Animal Health A/S, Fredensborg, Denmark; or Orion pharma, Espoo, Finland]). Pupils were dilated with a drop of 1% tropicamide (Mydriacyl; Alcon Nordic A/S, Copenhagen, Denmark), or a mixture of metaoxedrin-tropicamide drops (Oftan Tropicamide, 5mg/ml, Oftan metaoxedrin 100 mg/ml; Santen Oy, Tampere, Finland; mixed in a 1:5 ratio; for ERG/VEP recordings) and eyes lubricated with carbomer eye gel (Viscotears 2 mg/mL, Alcon Nordic). KM anesthesia was reversed by atipamezole 0.5–1 mg/kg (Antisedan, Orion Pharma, Copenhagen, Denmark).

### Intravitreal injections

Intravitreal injections were performed under an OPMI 1 FR PRO Surgical microscope (Zeiss, Jena, Germany) or Leica S9D Stereo microscope (Leica Microsystems, Wetzlar, Germany). A 30G disposable needle was used to puncture the sclera near the limbus, and a 33G or a 34G blunt-ended needle connected to a Hamilton syringe (Hamilton Company, Reno, NV, USA) was then inserted into the opening followed by injection of 1 µL hg-C_3_N_4_ NPs (1 mg/mL), red fluorescent silica particles (sicastar®-Red F, cat# 40-00-101, Micromod, Rostock, Germany), SiO_2_ (silica) particles, or PBS buffer solution. In the study with rd10 mice, Red F and SiO_2_ particles were used as controls instead of PBS to more closely mimic the physicochemical nature of hg-C_3_N_4_ NPs. For the bioavailability and safety study, both eyes of three WT mice were injected with NP solution and one mouse with buffer solution. Seven rd10 mice received hg-C_3_N_4_ NPs, four mice Red F particles, and three mice received SiO_2_ particles bilaterally.

### Fundus imaging and optical coherence tomography

Non-invasive fundus imaging was performed at baseline and 14 days following intravitreal injections in the safety study conducted in C57BL/6JRj mice using the Micron IV image-guided OCT 2 system (Phoenix Research Laboratories, Pleasanton, CA, USA). Similar imaging was performed in rd10 mice immediately after the intravitreal injection, and again 1 month after. Cross-sectional OCT B-scans and full volume 3D scans (only in some rd10 mice) were acquired centered on the optic disc. The B-scan images were segmented manually after loading the OCT scans into InSight software (Phoenix Research Laboratories). The coordinates were exported as .csv files and loaded into R (version R-4.3.0). The measurements were recentered around the optic nerve and the distance from the outermost border was calculated. Measurements less than 100 µm and further than 600 µm from the optic disc center were excluded. Mean total retinal thickness, mean NGI-thickness (combined thickness of retinal nerve fiber layer, retinal ganglion cell layer, and inner plexiform layer), mean combined thickness of the inner nuclear layer and outer plexiform layer, and mean outer nuclear layer thickness were calculated for each eye.

### Bulk RNA-seq library preparation and RNA-seq

Mice were intravitreally injected with 1 µL hg-C_3_N_4_ NPs (1 mg/mL) or PBS. Fourteen days post-injection retinas were dissected from enucleated eyes (n = 5) and total RNA purified from homogenized tissue using a mini hand-held homogenizer (Labdex, London, UK) and RNease total mini kit (Qiagen). Next, RNA was quantified and quality-checked prior to library preparation. Reverse transcription (RT) was performed using barcoded oligo(dT) primers to selectively capture polyadenylated mRNAs and introduce unique sample identifiers. RNA was mixed with barcoded oligo(dT) primers and dNTPs, followed by annealing at 65°C. First-strand cDNA synthesis was carried out using Maxima H Minus Reverse Transcriptase in the presence of a template-switching oligo (TSO) and PEG8000 to enhance strand-switching efficiency. The RT program included a 90-min incubation at 42 °C followed by 10 cycles alternating between 50 °C and 42 °C, with a final inactivation at 85 °C.

The resulting barcoded first-strand cDNAs were pooled and purified with 0.6× SPRIselect beads. Purified products were subjected to full-length cDNA amplification using KAPA HiFi HotStart ReadyMix with forward and reverse primers complementary to the barcode adapter sequences. The amplification program comprised initial denaturation at 95 °C for 3 min, followed by 12 cycles of 98 °C for 20 s, 57 °C for 15 s, and 72 °C for 2 min, with a final extension at 72 °C for 5 min. Amplified cDNA was size-selected and purified with 0.6× SPRIselect beads.

For library construction, amplified cDNA was processed using the Illumina-compatible library preparation workflow with library construction kit (10X Genomics). Briefly, cDNA underwent fragmentation, end-repair, and A-tailing, followed by adaptor ligation with Illumina-compatible dual index. Libraries were then purified by double-sided size selection using SPRIselect beads. Final library concentrations were measured by Qubit dsDNA HS assay, and quality were confirmed on an Agilent Bioanalyzer High Sensitivity DNA assay. Libraries were sequenced on an Illumina NovaSeqX plus platform.

### Retinal sectioning and flat mounting

Following sacrifice of the animals at day 14 post-injection eyes were enucleated and fixed in 4% stabilized formaldehyde buffer (VWR, Søborg, Denmark) overnight at 4 °C. One eye from each animal was used for flat mounting: The anterior segment and lens were removed and the neuroretina was carefully peeled off the retinal pigment epithelium (RPE)/choroid and immersed in ice-cold PBS buffer before proceeding with immunostaining. The other eye was processed for paraffin embedding and sectioning at the Department of Pathology, Aarhus University Hospital.

### Electroretinography

Overnight dark-adapted mice were anesthetized under dim red-light observation with ketamine (60 mg/kg) and medetomidine (0.4 mg/kg). Electroretinography (ERG) recordings were performed as previously described ^98^. Pupils were dilated with applications of mydriatic eye drops (Oftan Tropicamid, 5 mg/mL, Oftan Metaoxedrin 100 mg/ml; Santen Oy, Tampere, Finland; mixed in a 1:5 ratio). Corneal protection and conductivity were ensured using carbomer-based eye lubricant (Viscotears, Bausch & Lomb Nordic AB, Stockholm, Sweden). Mice were maintained at 37°C on a heating pad throughout the procedure. Scotopic ERG responses were recorded using a Diagnosys Espion E3 system (Espion E3 console, Diagnosys LLC, Lowell, MA) with silver-wire corneal electrodes and subdermal reference in the snout and ground electrode in the lower back. Both eyes were stimulated simultaneously with monochromatic green light across 12 increasing intensity steps (0.00001 to 30 cd·s/m²), with inter-stimulus intervals ranging from 1 to 60 seconds and 2–25 sweeps per step. Signals were sampled at 2 kHz and filtered (1–300 Hz). Only the response for the strongest flash (30 cd·s/m^2^) was used for final analysis.

### Light-dark box test

The light-dark box (LDB) contains a plastic chamber (44 x 18 x 21 cm) divided into two equal compartments by a divider, which had a small passage that allowed animals to move freely between the compartments. One compartment is brightly lit (∼900 lux) with a lamp above the arena, whereas the other compartment is darkened (∼5 lux) getting light mostly from the lit side through the passage. Two days before the experimental day, all animals were allowed to familiarize themselves to the test environment for 3 minutes. In the actual experiment, each animal was placed initially in the lit compartment and was allowed to freely move in the apparatus (between the chambers) for 5 minutes in a calm and silent environment. Animal behavior was recorded using a digital camera positioned above the light zone. A researcher blinded as to the treatments analyzed the total time spent in the dark compartment.

### Cortical visual evoked potential recordings and analysis

Carprofen (20 mg/kg s.c; Rimadyl Vet 50 mg/ml, Zoetis, Helsinki, Finland) was prophylactically administered as a pain killer. Mice were initially anesthetized with isoflurane in an induction chamber (concentration 4.5 %) and then transferred to stereotaxic frame (Kopf Instruments, Tujunga, CA, USA) where isoflurane was administered through a nose mask. At this point, also urethane anesthesia (2 g/kg, s.c.) was induced and the pupils were dilated with a mixture of metaoxedrin-tropicamide drops (5 mg/ml and 100 mg/ml, respectively, mixed in a 1:5 ratio). During surgical procedures, isoflurane concentration was gradually lowered from 1.5 % to 0.6 % until finally stopped (leaving animal to be anesthetized with urethane only) when surgical operations were finalized. The surgery was performed as previously described ^99^. The skin over the scalp was locally anesthetized with lidocaine (20 mg/ml, Xylocaine, Aspen Pharma Trading Limited, Irland). Thereafter the scalp was opened with an incision and the skull was cleaned. Holes for electrodes were drilled with a dental drill leaving the dura mater intact. Two stainless steel mini screw (shank diameter 1 mm) were attached on the holes that were drilled bilaterally above the binocular primary visual cortex (medial/lateral: ± 3 mm from lamda), and one on frontal bone (AP: 2 mm; ML: 1 mm) which served as the reference electrodes. The visual evoked potentials (VEPs) were recorded with a Diagnosys Celeris rodent ERG device (Diagnosys LLC, Lowell, MA). The eyes were stimulated one at a time with an ascending (log unit intervals) stimulus intensity series as follows: Step 1, a 4 ms flash at 0.05 cd·s/m² was delivered with 50 to 100 repetitions (the number of flash repetitions varied depending on the background EEG signal strength and waveform) and a 1-s inter-stimulus interval (ISI). Step 2 used a stimulus intensity of 0.5 cd·s/m², repeated 40 to 80 times with a 1.5-second ISI. In Step 3, the intensity was increased to 5.0 cd·s/m², with 100 to 200 repetitions and a 5-second ISI. Step 4 involved stimulation at 50 cd·s/m², repeated 80 to 160 times with a 7.5-second ISI. Finally, Step 5 used the highest intensity of 500 cd·s/m², with 40 to 120 repetitions and a 7.5-second ISI. Sweeps were manually inspected offline and consistent and clean sweeps averaged for waveform analysis. The EEG signal was collected at 2 kHz and bandpass-filtered between 0.25–300 Hz.

To enable unbiased identification of eyes exhibiting cortical VEP responses, we implemented an objective responder classification pipeline based on four quantitative waveform features. For each stimulus intensity, VEP responses were evaluated within a post-stimulus window of 80–220 ms, using the −50 to 0 ms interval as baseline. A response was considered “responder-like” if it satisfied at least three of the following four criteria: (1) Positive-polarity signal-to-noise ratio (SNR ≥ 4), computed as the maximum positive deflection relative to the baseline standard deviation. (2) Positive-energy ratio ≥ 0.6, defined as the proportion of signal energy above baseline within 80–220 ms. (3) Peak latency between 80 and 220 ms after stimulus onset. (4) Waveform template correlation (Pearson r ≥ 0.4), where for each intensity, a template VEP waveform was generated as the leave-one-out average of all other eyes recorded at the same intensity. This approach avoids bias toward any treatment group and provides an intensity-specific representation of VEP shape against which each candidate waveform is compared. The analysis was implemented over four stimulus intensities (0.5, 5, 50, and 500 cd·s/m²). Eyes were classified as VEP responders if three or more intensities met the responder-like criteria. Responder fractions were quantified for each treatment group, and Wilson binomial 95 % confidence intervals were computed. Differences in responder probability between groups were expressed as risk differences with Newcombe 95 % confidence intervals. All waveform processing, responder classification, and statistical computations were performed in Python

### Porcine retina dissection

Eyes from Danish Land Race pigs (*Sus domesticus*) were collected at a local slaughterhouse (Danish Crown, Horsens, Denmark) immediately after the animals had been anesthetized with carbon dioxide and killed by exsanguination. The eyes were transported to Aarhus University Hospital (AUH) laboratory in physiological saline solution (PSS) at 4 °C, and the time from the collection of the eyes to the commencement of the dissection procedure never exceeded 5 h. Dissection was performed in 4 °C PSS0.0 as follows: each eye was bisected at the equator with a double-edged blade, and the anterior segment was removed. The posterior segment containing the optic disc was placed under a stereo microscope, the vitreous body was removed. A few mm from the optic disc a segment of approximately 3 x 3 mm^2^ neuroretinal tissue was cut out using a self-locking chisel blade handle (VWR International, Herlev, Denmark) equipped with a 30 microblade (BD Beaver, D.J. Instruments, Billerica, USA). Chemical Solutions for storage and transportation: PSS with the following composition (in mM): 118 NaCl, 4.8 KCl, 1.14 MgSO_4_, 25 NaHCO_3_, 5 Hepes, 1.5 CaCl_2_, 5.5 glucose.

### MEA electrophysiology

A micro-electrode array (MEA) with 32 channels from Blackrock Microsystems was used to measure electrical signals from the retinal tissue. The MEA was connected via an Omnetics connector to a Blackrock Microsystems Cereplex Direct dedicated data acquisition. Data was sampled at 30 kHz, high pass filtered at 250 Hz and further filtered for 50 Hz line noise by a built-in Blackrock Line Noise Cancelation filter (filtering 50 Hz, 1 second, soft synchronization). For all recordings except those used for dose–response analysis, a 50 Hz notch filter was additionally applied in MATLAB to attenuate residual line-noise artefacts. Data was analyzed using custom Matlab software and python scripts. All electrophysiological experiments were carried out inside a Faraday cage.

After placing the retinal tissue on the MEA with the retinal ganglion cells facing upwards, 5 µg of hg-C_3_N_4_ NPs dispersed in 20 µL of PBS we added to tissue and allowed to settle for around 15 min. Afterwards, light stimulation was applied using a 450 nm homemade LED system with 1 ms pulses with one pulse per second (1 Hz). The recording length was at least 30 s. The resulting whole data trace for each of the 32 channels were segmented into traces of 1 s duration (minimum 30 traces), initiating at the stimulus artefact. These traces were then averaged in order to improve the signal/noise ratio. Firing-rate analyses were performed on non-averaged traces to avoid emphasizing residual line-noise artefacts, and spikes were detected using a median absolute deviation (MAD)-based thresholding approach. Briefly, spikes were detected on each channel using a threshold set at 4.0 × MAD of the baseline noise, calculated for each trace independently. Firing frequency was quantified by counting detected spikes within a 1–100 ms window following light onset for each stimulus. All spike detection and firing-rate analyses were implemented using custom Python scripts.

Chemical Solutions for experiments: PSS1.6 with the following composition (in mM): 119 NaCl, 4.7 KCl, 1.17 MgSO_4_, 25 NaHCO_3_, 5 Hepes, 1.6 CaCl_2_, 5.5 glucose, 1.18 KH_2_PO_4_, 0.026 EDTA. PSS0.0 refers to PSS1.6 where CaCl_2_ has been omitted. PSS1.6 was heated to 37°C and oxygenated prior to all experiments in order to ensure a physiological environment during electrical measurements.

Tetrodotoxin (TTX) citrate was obtained from Hello Bio Reagents (Bristol, United Kingdom, HB1035). TTX solutions were prepared by creating a stock solution of TTX in PSS1.6 of 1.0 M concentration. From this solution 1 mL aliquots of 2.25 mM concentration were created for use in future experiments. All TTX solutions were stored at -20°C between experiments. For TTX control experiments, a volume of 50 µL of the TTX solution was applied directly to the tissue sample.

### NP uptake and *ex vivo* culture of porcine retina

Porcine eyes were obtained from the slaughterhouse as described previously. Periocular tissue was removed, and the eye was disinfected in a 10% w/v iodine solution (polyvinylpyrrolidone-iodine complex [Thermo Fisher Scientific, Roskilde, Denmark] in PBS). The anterior segment of the eye was cut off and blunt forceps were used to remove the vitreous. The remaining tissue was quartered and for each a 6 mm punch out of the retina was carefully transferred to a well with 1:1 DMEM/F12 (both Euroclone, Pero, MI, Italy) medium supplemented with 2% B-27 (Thermo Fisher Scientific), 1 µg/mL streptomycin, 1 U/mL penicillin, and 2 mM L-glutamine (all Euroclone). Media was aspirated and 30 µL NP solution (2 mg/mL) was added directly onto the retinas and incubated for 30 minutes. Afterwards media was added, and retinas incubated for 24 hours.

### Immunostaining of retinal flat mounts and sections

Porcine retinas were fixed for 6 hours at RT in 4% stabilized formaldehyde buffer (VWR), washed three times in PBS, and blocked and permeabilized overnight at 4°C in retina blocking buffer (RBB) containing PBS with 1% BSA (VWR) and 0.5% Triton X100 (Sigma, Søborg, Denmark). Next, retinas were incubated with rabbit anti-RbPMS (RNA binding protein with multiple splicing, RGC marker) (1.6 mg/mL; NBP2-20112, Novus Biologicals, Littleton, ON, Canada) diluted 1:500 for 48 hours at 4 °C. Following wash in wash buffer (PBS with 0.5% Triton X100) they were incubated with Alexa-568 Donkey anti-rabbit (2 mg/mL; A10042, Thermo Fisher Scientific) diluted 1:400 overnight at 4 °C). Retinas were then washed and counterstained with Draq5 (5 mM; 62251, Thermo Fisher Scientific) diluted 1:1000 in PBS for 10 minutes. Finally, retinas were washed and transferred to glass slides, where they were mounted with the RGC layer facing upwards using Fluoromount-G mounting medium (Thermo Fisher Scientific).

Mouse flat mounts were blocked and permeabilized overnight at 4 °C in RBB before being incubated with rabbit anti-RbPMS (1.6 mg/mL; NBP2-20112, Novus Biologicals) diluted 1:400 in RBB overnight at 4 °C. Next, the retinas were washed and incubated with Alexa-568 goat anti-rabbit (2 mg/mL; A11011, Thermo Fisher Scientific) diluted 1:400 in RBB overnight at 4°C. The flat mounts were washed and counterstained with Draq5 (5 mM; 62251, Thermo Fisher Scientific) diluted 1:1000 in wash buffer before they were transferred to glass slides and mounted with the RGC layer facing upwards using Fluoromount-G mounting medium (Thermo Fisher Scientific).

Mouse retinal sections were deparaffinized in xylene overnight before rehydration with graded ethanol washes. Heat-induced epitope retrieval was performed in citrate buffer pH 6 for RbPMS and Tris-EDTA buffer (10 mM Tris and 1 mM EDTA, Sigma) pH 9 for Iba1. Antigen retrieval was performed with proteinase K (20 µg/ml in PBS; Promega, Madison, Wisconsin, USA) for GFAP. Sections were blocked with 3% BSA and 1% Triton X100 in PBS for 1 hour before incubation with anti-RbPMS (1 mg/mL; NBP2-20112, Novus Biologicals) diluted 1:400, anti-GFAP (3.2 mg/mL; Z0334, Dako, Glostrup, Denmark) diluted 1:500, or anti-Iba1 (0.5 mg/mL; 019-19741, FUJIFILM Wako Pure Chemical Corporation, Osaka, Japan) diluted 1: 700 in PBS with 1% BSA overnight at 4°C. The next day, sections were washed and incubated with Alexa-488 goat anti-rabbit (2 mg/mL; A11008, Thermo Fisher Scientific) diluted 1:400 in PBS with 1% BSA for 30 minutes. The sections were washed and counterstained with Draq5 (5 mM; 62251, Thermo Fisher Scientific) diluted 1:1000 in PBS and mounted with Fluoromount-G mounting medium (Thermo Fisher Scientific) on glass slides.

For post-mortem evaluation after ERG, LDB, and VEP tests, mouse eyes were fixed overnight in Hartmańs fixative (H0290, Sigma-Aldrich) and thereafter transferred to 70 % ethanol. Eyes were embedded in paraffin and sectioned at 5 µm. For standard hematoxylin-eosin histology staining, the sections were heated at 50 °C for 30 min, then deparaffinized in xylene (2×5 min), and an ethanol series (99.9 % 2×2 min, 94 % 2×2 min, 70% 1×5 min, and 50 % 1×5 min), and rehydrated in water (1×20 sec). For immunostaining, the sections were deparaffinized in xylene (3×5 min) and ethanol series (99.9 % 2×5 min, 94 % 1×5 min, 70% 1×5 min and 50 % 1×5 min), rehydrated in water (1×1 min) and washed in phosphate buffered saline (PBS, pH 7.4,), after that, the sections were blocked with 5 % normal donkey serum (NDS), prepared in PBS, for 1 hour. Next, the sections were incubated in rabbit anti M-opsin antibody (NB110-74730, Novus Biologicals, dilution 1:250), prepared to 5 % NDS-PBS-0.1 % Triton X solution, with mild orbital shaking overnight at 4 °C. After that, the sections were washed for 3×5 mins in PBS 0.1% triton (PBST) and incubated with a fluorescent secondary antibody (CoraLite donkey anti-rabbit 488nm, SA00013-6, ProteinTech, dilution factor 1:500) for 2 hours in the dark, at room temperature with mild shaking. Each slide contained at least one negative control sample. Finally, the slides were washed again for 3×5 min in PBST 0.1 % and 1×5 min in PBS to ensure removal of residual detergent from the glass surface. The sections were dried and subsequently mounted using Fluoroshield mounting medium with DAPI (ab104139, Abcam).

### Imaging of retinal sections

Image acquisition and analyses were performed at the Bioimaging Core Facility, Health, Aarhus University, Denmark. Retinal sections were imaged with the Olympus VS120 upright widefield fluorescence microscope (Olympus, Tokyo, Japan) equipped with Spectra X 7IR LED multi-spectral light engine and Semrock pentafilter (DAPI/FITC/Cy3/Cy5/Cy7 Penta LED HC Filter Set #F68-050) with Hamamatsu ORCA-FLASH4.0 V2 (QE 82%) camera (Hamamatsu, Shizuoka, Japan). Overview images were taken with the Olympus UPlanSApo 20x/0.75 air objective and associated VS-ASW imaging software. Iba1 and GFAP IF on retinal sections were imaged with the Olympus BX63 upright widefield fluorescence microscope equipped with a CoolLED *p*E300^ultra^ illumination module and dedicated filters optimized for the DAPI (λ_ex_ = 360-370 nm, λ_em_ = 420-460 nm), for the Alexa Fluor 488 (λ_ex_ = 465-495 nm, λ_em_ = 515-555 nm), and for the Cy5 (λ_ex_ = 590-650 nm, λ_em_ = 663-737 nm) channels with Andor Zyla 5.5 sCMOS (QE 60%) camera (Andor Technology, Belfast, Northern Ireland). 40x images were taken with the Olympus UPlanFl 40x/0.75 air objective and associated CellSens imaging software.

Retinal sections and flat mounts were imaged with a Zeiss LSM 800 confocal laser scanning microscope (Zeiss, Jena, Germany) using a Plan-Apochromat 63x/1.4 oil objective. Excitation and emission wavelengths were λ_ex_ = 405 nm and λ_em_ = 400-605 nm for the DAPI channel, λ_ex_ = 488 nm and λ_em_ = 400-650 nm for Alexa Fluor 488, λ_ex_ = 561 and λem = 400-650 for Alexa Fluor 568, and λ_ex_ = 640 and λ_em_ = 650-700 for the Cy5 channel.

Images were captured with fixed settings and processed similarly. Display settings are similar and fluorescence intensities are comparable among images in the same figure. HE histology and M-opsin immunostained specimens were imaged using an Olympus APEXVIEW APX100 microscope equipped with an Olympus U-LGPS light source. All zoomed-in images were acquired from the central region of the retina.

### Statistical analysis

Statistical analyses were performed using GraphPad Prism (versions 9.3.1 or 10.4.1) or R (version 4.3.0). Statistical significance was assessed using unpaired two-tailed t-tests or one-way ANOVA and Turkey’s test for multiple comparison unless otherwise stated. P-values are indicated in the figures where p<0.0001 is denoted as ****.

### Reporting Summary

Further information on research design is available in the Nature Research Reporting Summary linked to this article.

## Data availability

The main data supporting the results in this study are available within the paper and its Supplementary Information. Source data for the figures will be provided with this paper. All data generated in this study will be available from Zenodo link.

## Acknowledgments

M. Chen gratefully acknowledges the Carlsberg Foundation (CF19-0300) and Novo Nordisk Foundation (NNF22OC0080508), Lundbeck Foundation (R400-2022-1232), Marie Skłodowska-Curie Actions programme (ENSIGN, grant number 101086226 and L4DNANO, grant number 101086227), for supporting this work. B. Tian acknowledges the support from the Ziyi-Zhong gift. T. Bek, J. G. Madsen and R. S. Davidsen gratefully acknowledge the Novo Nordisk Foundation (NNF20OC0064628,). T. Bek and T.J. Corydon acknowledges the APT Holding. H. Leinonen acknowledges support from the Research Council of Finland, Business Finland, Emil Aaltonen Foundation, Sigrid Jusélius Foundation, Päivikki and Sakari Sohlberg Foundation, Finnish Eye and Tissue Bank Foundation, Retina Registered Association (Finland), and Sokeain Ystävät/De Blindas Vänner Registered Association. T.J. Corydon acknowledges the VELUX Foundation (Grant No. 00038189), and the Independent Research Fund Denmark (Grant No. 2034-00036B). A.L. Askou gratefully acknowledges The Danish Eye Research Foundation. T.S. Jakobsen acknowledges Fight for Sight, Denmark and the Synoptik Foundation. NSH acknowledges funding from The Novo Nordisk foundation challenge center for Optimised Oligo Escape and Control of Disease (NNF23OC0081287), and The Center for 4D Cellular Dynamics (NNF22OC0075851) Lundbeck foundation (R453-2024- 589 359) Villum Experiment grant (40801), and the Swiss National Science Foundation 598 (310030M_204518). NSH acknowledges funding from The Novo Nordisk foundation challenge center for Optimised Oligo Escape and Control of Disease (NNF23OC0081287), and The Center for 4D Cellular Dynamics (NNF22OC0075851) Lundbeck foundation (R453-2024- 589 359) Villum Experiment grant (40801), and the Swiss National Science Foundation 598 (310030M_204518). Fig. 1b-d and 4a were partly generated using Servier Medical Art, provided by Servier, licensed under a Creative Commons Attribution 3.0 unported license. The authors acknowledge the Bioimaging Core Facility, Department of Biomedicine, Aarhus University, Denmark, for the use of equipment and support. The authors thank Tina Hindkjær for excellent technical assistance.

## Funding

Carlsberg Foundation (CF19-0300) Ziyi-Zhong gift

Novo Nordisk Foundation (NNF20OC0064628) Novo Nordisk Foundation (NNF23OC0081287) Novo Nordisk Foundation (NNF22OC0075851)

Swiss national science foundation 310030M_204518 APT Holding

VELUX Foundation (00038189) and 40801 Independent Research Fund Denmark (2034-00036B) Fight for Sight

The Danish Eye Research Foundation Synoptik Foundation

Lundbeck foundation (R453-2024- 589 359Swiss National Science Foundation 598 (310030M_204518

## Author contributions

Conceptualization: MC, BT

Methodology: MC, BT, HL, CAM, KKK, JZ, PL, LM, WL, TSJ, ALA, TJC

Investigation: YZ, JZ, PL, SVB, GB, LM, JGM, KKK, TSJ, ACJ, AA, AK,GH, TSJ,

Visualization: MC, HL, KKK, YZ, TSJ, ALA

Supervision: MC, BT, TB, MD, RSD, TJC, NSH, ALA

Writing-original draft: CAM, KKK, CM, BT, TB, JGM, RSD, MD, TSJ, ALA, TJC

