## supplementary file for "Biomimetic graphitic carbon nitride nanoparticles enable multiscale biomodulation"

### Biomimic opto-nanobiointerface enables multiscale biomodulation

**Affiliations**


**This Supplementary file includes:**

Fig. S1 to S7

**Other Supplementary Materials for this manuscript include the following:**

Videos S1 to S12

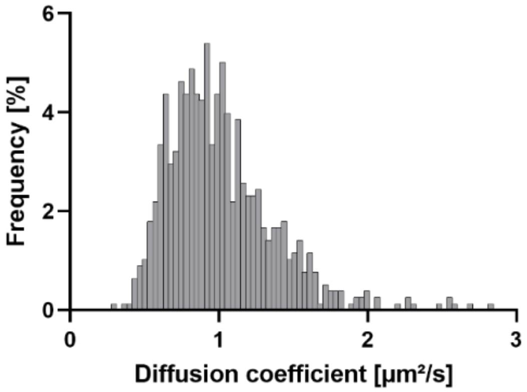


**Fig. S1.**

Diffusion coefficient distribution of hg-$C_3N_4$ derived from particle tracking analysis.

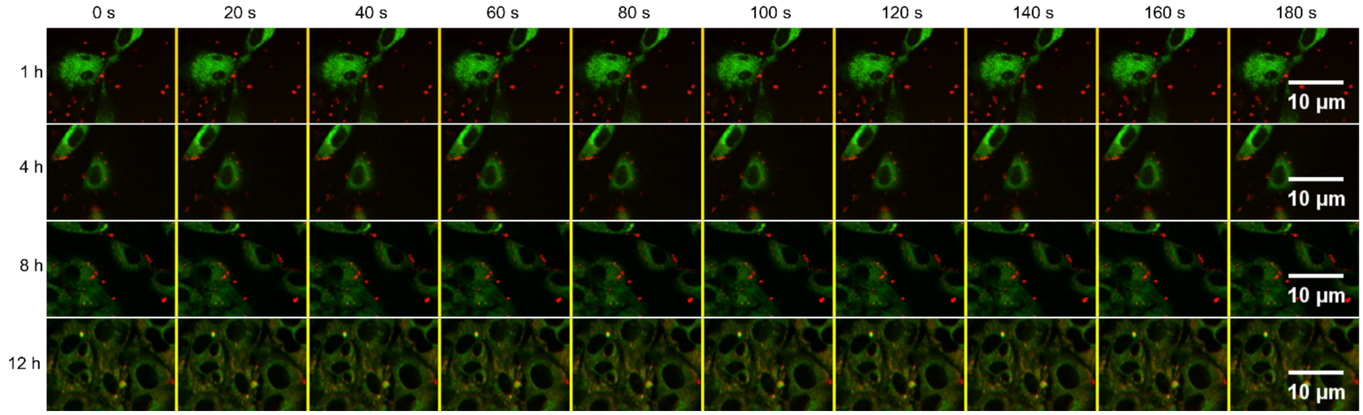


**Fig. S2.**

**Images acquired with SPT used for colocalization study.** NIH/3T3 fibroblasts where lysosomes are stained with LysoTracker Deep Red (green) and hg-$C_3N_4$ NP excited with 488 nm shown in red. The cells were imaged at 1h, 4h, 8h, and 12h with 1 frame per 20 seconds for 3 min.

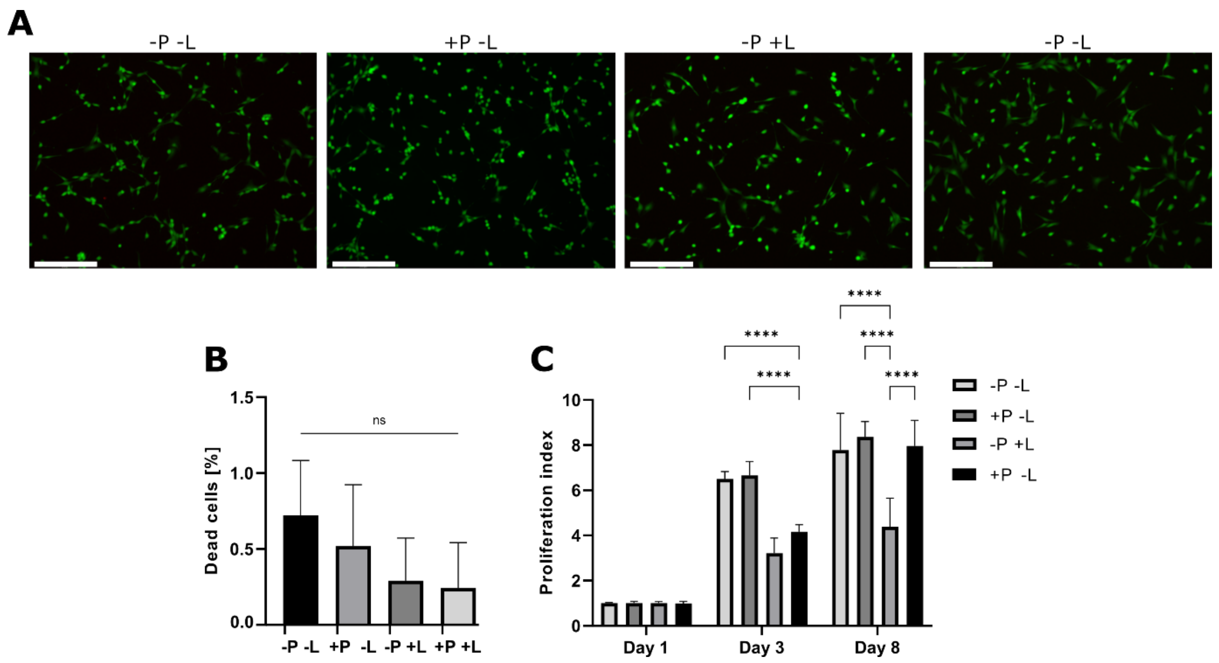


**Fig. S3.**

**Cytocompatibility of hg-$C_3N_4$. (A)** Live dead staining of NIH/3T3 fibroblasts with **(B)** percentage of dead cells based on live dead staining and **(C)** CCK-8 proliferation assay of NIH/3T3 cells cultured with 2.0 μg/cm$^2$ hg-$C_3N_4$ (+P), cultured without hg-$C_3N_4$ (-P), light treated (+L), and non-light treated (-L). Light-treated groups were irradiated with a 450 nm blue light (75 mW/cm$^2$, 1 Hz, 100 ms pulse) for 10 minutes bidaily. Statistical analysis was performed with one-way ANOVA and Tukey's test for multiple comparisons where $p<0.0001$ is denoted as **** with n=7 and ns=no significance ($p>0.05$)

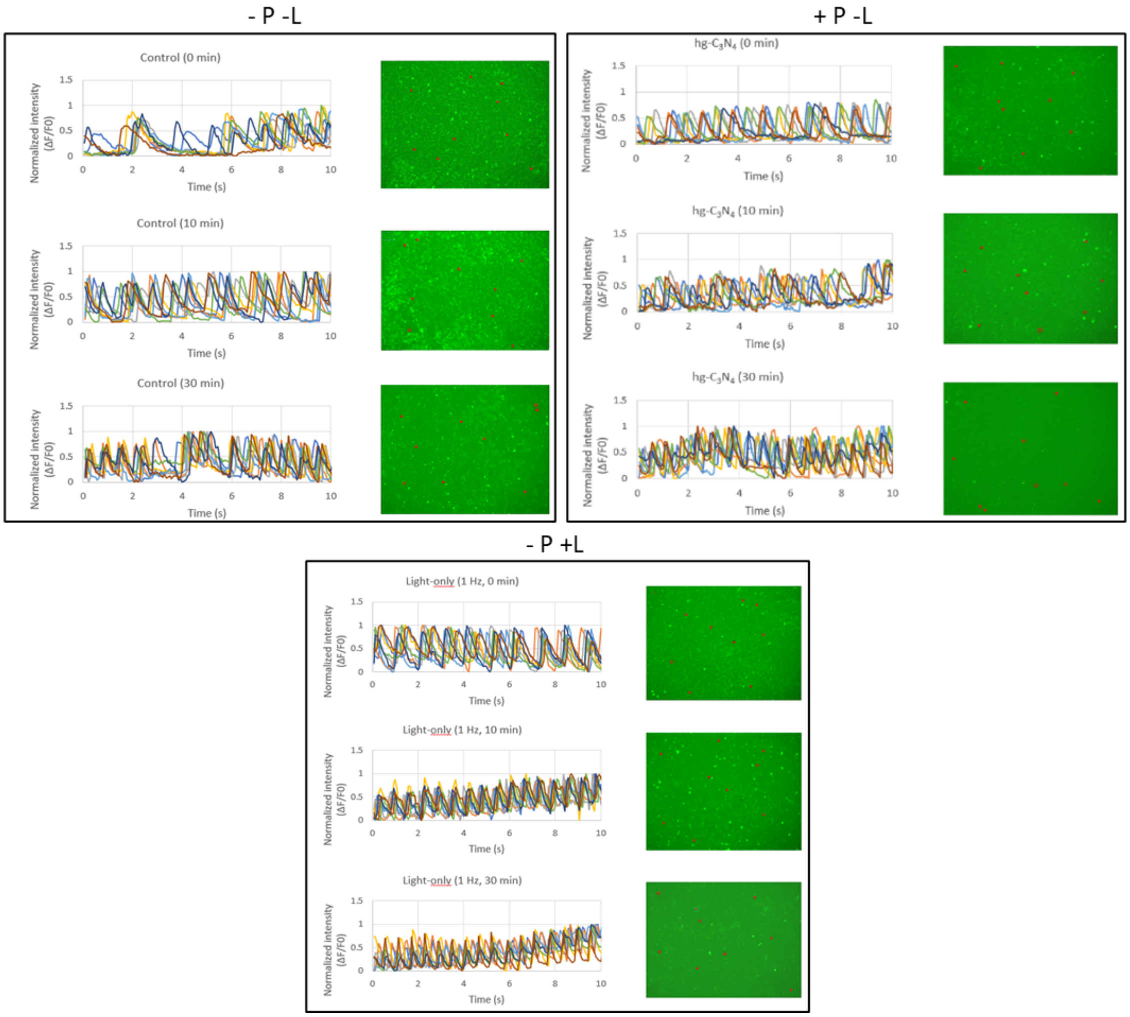


**Fig. S4.**

**Control groups for HL-1 cardiomyocyte pacing**. HL-1 cells stained with fluo-4 calcium stain incubated with or without 2.0 μg/cm$^2$ hg-$C_3N_4$ particles (+P; -P) and with or without photostimulation photostimulation (+L; -L) at 0 min, 10 min, and 30 min. Red circles indicate ROIs for $\Delta F/F_0$ plots. $\Delta F/F_0$ plots show 8 different ROIs showing beating frequency of HL-1 cells. Light source was 10 mW/cm$^2$ of 450 nm with pulse width of 100 ms and 1 Hz frequency.

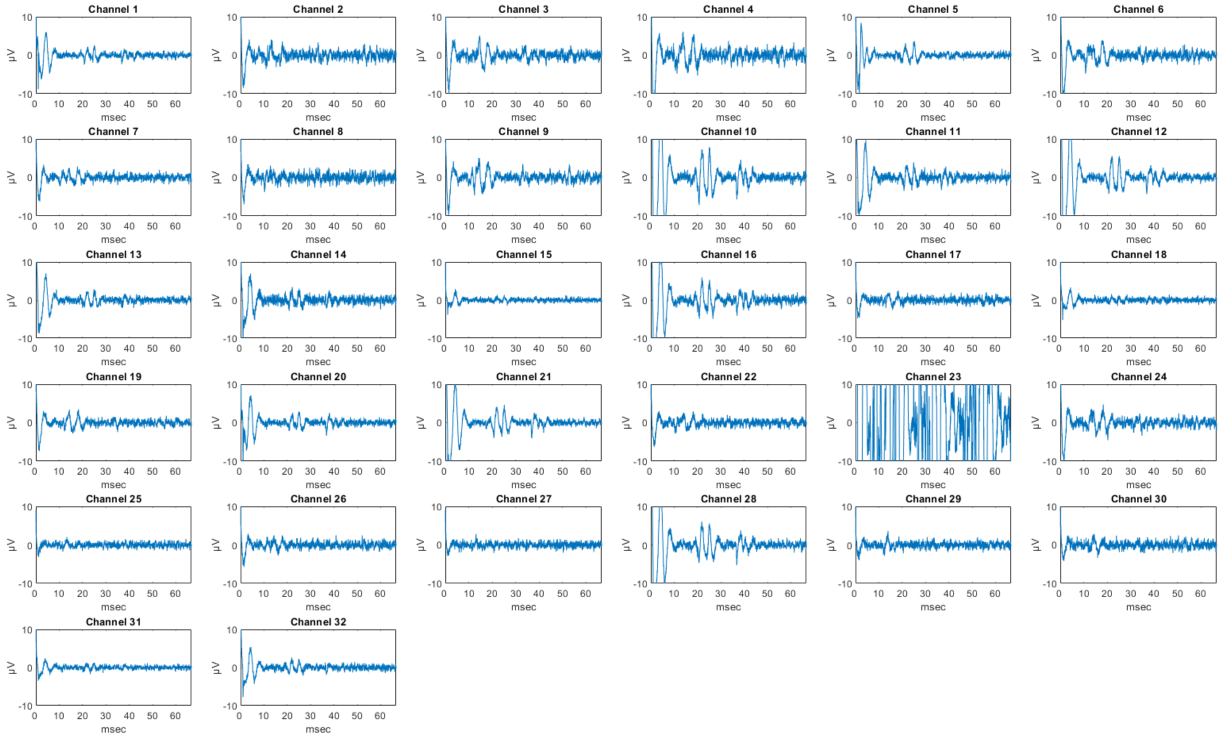


**Fig. S5**

Stimulation response data of all 32 MEA channels before applying particles with LED stimulation of 1 msec duration. Stimulus artefact and evoked light responses can be seen on all channels, except channel 23 which malfunctioned.

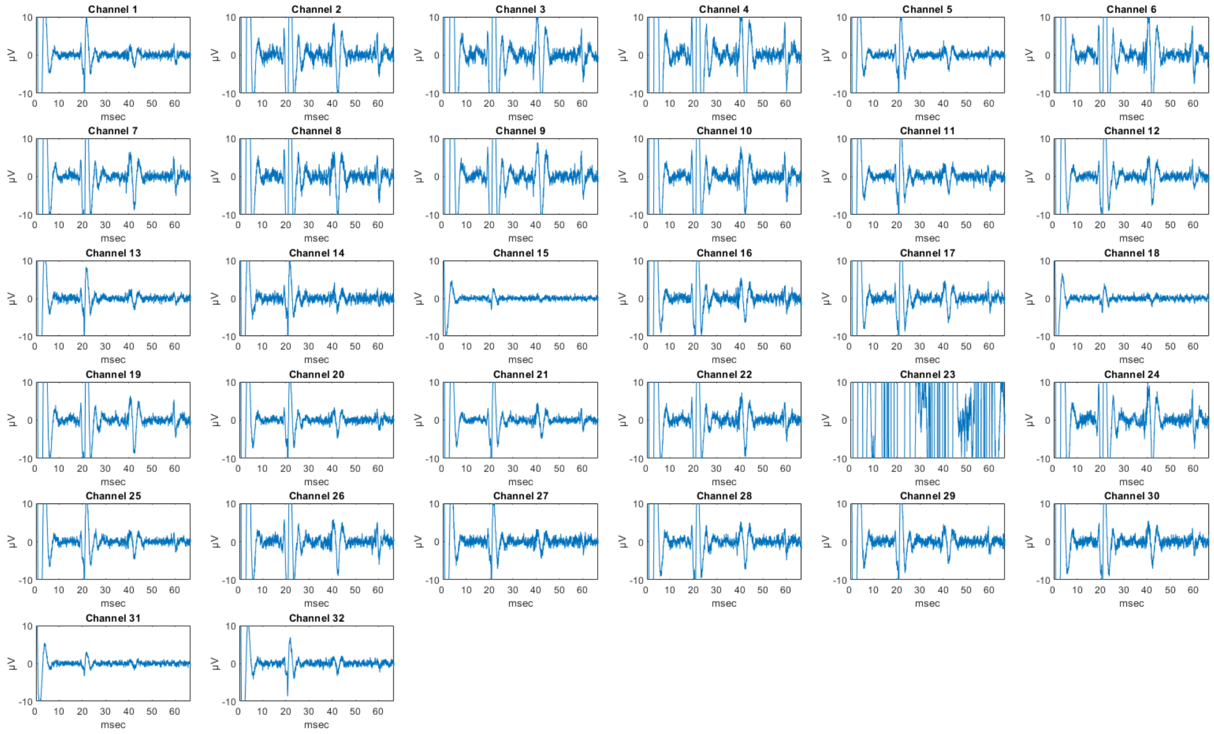


**Fig. S6**

Stimulation response data of all 32 MEA channels after applying particles with LED stimulation of 1 msec duration. Stimulus artefact and evoked light responses can be seen on all channels, except channel 23 which malfunctioned. All evoked potentials display larger peak-peak amplitude than before particle application.

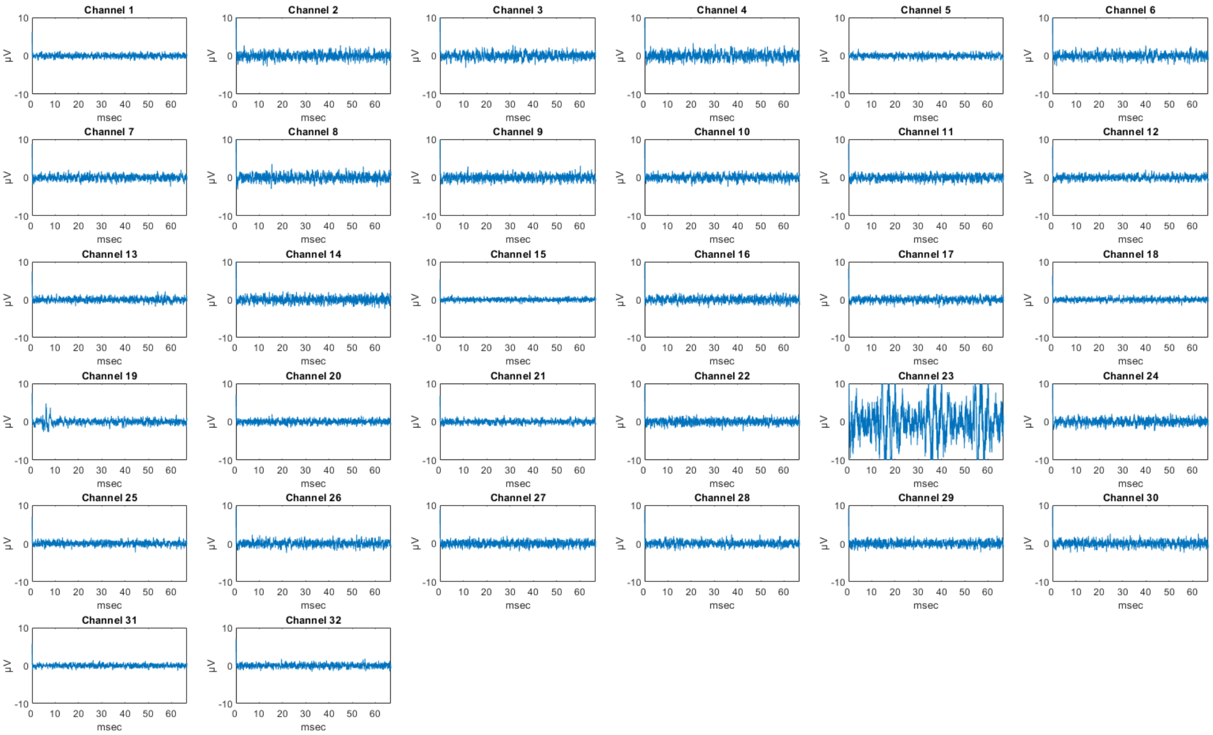

**Fig. S7**

Stimulation response data of all 32 MEA channels after applying TTX with LED stimulation of 1 msec duration. Stimulus artefact can still be seen after TTX application, but no clear evoked potentials are visible on any channel.

### Video descriptions

**S1 hg-$C_3N_4$ Internalization**

Internalization of hg-$C_3N_4$ by single NIH/3T3 fibroblast cell

**S2 Laser stimulation of NIH/3T3 fibroblasts without hg-$C_3N_4$ NPs**

Laser stimulation of 4 NIH/3T3 fibroblast cells as non-particle control with 200 ms laser pulse, 473 nm, spot size 3 µm, and 8.8 mW/µm$^2$. Green fluorescence is calcium staining.

**S3 Laser stimulation of NIH/3T3 fibroblast cell with hg-$C_3N_4$ NPs**

Laser stimulation of internalized hg-$C_3N_4$ NPs of single NIH/3T3 fibroblast cell with 200 ms laser pulse, 473 nm, spot size 3 µm, and 8.8 mW/µm$^2$. Green fluorescence is calcium staining and hg-$C_3N_4$ NPs. Red arrow indicates location of laser stimulation.

**S4 Laser stimulation of NIH/3T3 fibroblast cell with hg-$C_3N_4$ NPs as $\Delta F/F_0$ video.**

$\Delta F/F_0$ video of laser stimulation of internalized hg-$C_3N_4$ NPs of single NIH/3T3 fibroblast cell with 200 ms laser pulse, 473 nm, spot size 3 µm, and 8.8 mW/µm$^2$. $\Delta F/F_0$ video is generated based on Supplementary video S3.

**S5 Laser stimulation of HELA cell with hg-$C_3N_4$ NPs**

Laser stimulation of internalized hg-$C_3N_4$ NPs of single NIH/3T3 fibroblast cell with 200 ms laser pulse, 473 nm, spot size 3 µm, and 8.8 mW/µm$^2$. Green fluorescence is calcium staining and hg-$C_3N_4$ NPs. Red arrow indicates location of laser stimulation.

**S6 Laser stimulation of HELA cell with hg-$C_3N_4$ NPs as $\Delta F/F_0$ video.**
Laser stimulation of internalized hg-$C_3N_4$ NPs of single NIH/3T3 fibroblast cell with 200 ms laser pulse, 473 nm, spot size 3 µm, and 8.8 mW/µm$^2$. $\Delta F/F_0$ video is generated based on Supplementary video S5.
**S7 Laser stimulation and signal propagation of CMs with hg-$C_3N_4$ NPs**
Laser stimulation of CM with hg-$C_3N_4$ NPs resulting in immediate intracellular calcium flux and beating in all 4 adjacent CMs. Stimulation with 200 ms laser pulse, 473 nm, spot size 3 µm, and 8.8 mW/µm$^2$. Green fluorescence is calcium staining and hg-$C_3N_4$ NPs. Red arrow indicates location of laser stimulation.
**S8 Laser stimulation and signal propagation of CFs with hg-$C_3N_4$ NPs**
Laser stimulation of CF with hg-$C_3N_4$ NPs resulting in immediate intracellular calcium flux and signal propagation to adjacent CF. Stimulation with 200 ms laser pulse, 473 nm, spot size 3 µm, and 8.8 mW/µm$^2$. Green fluorescence is calcium staining and hg-$C_3N_4$ NPs. Red arrow indicates location of laser stimulation.
**S9 Laser stimulation and signal propagation of CF to CM with hg-$C_3N_4$ NPs**
Laser stimulation of CF with hg-$C_3N_4$ NPs resulting in immediate intracellular calcium flux and signal propagation to adjacent CM. Stimulation with 200 ms laser pulse, 473 nm, spot size 3 µm, and 8.8 mW/µm$^2$. Green fluorescence is calcium staining and hg-$C_3N_4$ NPs. Red arrow indicates location of laser stimulation.
**S10 Pacing of HL1 CM cell calcium dynamics via LED stimulation of hg-$C_3N_4$ NPs: calcium dynamics before stimulation.**
HL1 cell calcium dynamics before application of LED stimulation. Green fluorescence is calcium staining.
**S11 Pacing of HL1 CM cell calcium dynamics via LED stimulation of hg-$C_3N_4$ NPs: after 10 min stimulation.**
HL1 cell calcium dynamics after application of 10 min pulsed LED stimulation. Green fluorescence is calcium staining. Light source was 10 mW/cm$^2$ of 450 nm with pulse width of 100 ms and 1 Hz frequency.
**S12 Pacing of HL1 CM cell calcium dynamics via LED stimulation of hg-$C_3N_4$ NPs: after 30 min stimulation.**
HL1 cell calcium dynamics after application of 30 min pulsed LED stimulation. Green fluorescence is calcium staining. Light source was 10 mW/cm$^2$ of 450 nm with pulse width of 100 ms and 1 Hz frequency.
